# Mitochondrial citrate carrier SLC25A1 is a dosage-dependent regulator of metabolic reprogramming and morphogenesis in the developing heart

**DOI:** 10.1101/2023.05.22.541833

**Authors:** Chiemela Ohanele, Jessica N. Peoples, Anja Karlstaedt, Joshua T. Geiger, Ashley D. Gayle, Nasab Ghazal, Fateemaa Sohani, Milton E. Brown, Michael E. Davis, George A. Porter, Victor Faundez, Jennifer Q. Kwong

**Affiliations:** Graduate Program in Biochemistry, Cell and Developmental Biology; Graduate Division of Biological and Biomedical Sciences; Emory University; Atlanta, GA, USA; Division of Pediatric Cardiology; Department of Pediatrics; Emory University School of Medicine; and Children’s Healthcare of Atlanta; Atlanta, GA, USA; Department of Cardiology; Smidt Heart Institute; Cedars-Sinai Medical Center; Los Angeles, CA, USA; Division of Vascular Surgery; University of Rochester Medical Center; Rochester, NY, USA; Wallace H. Coulter Department of Biomedical Engineering; Emory University School of Medicine; Atlanta, GA, USA; Department of Pediatrics; Division of Cardiology; University of Rochester Medical Center; Rochester, NY, USA; Department of Cell Biology; Emory University School of Medicine; Atlanta, GA, USA

**Keywords:** Mitochondria, heart, congenital heart disease, oxidative phosphorylation, metabolism, epigenetics

## Abstract

The developing mammalian heart undergoes an important metabolic shift from glycolysis toward mitochondrial oxidation, such that oxidative phosphorylation defects may present with cardiac abnormalities. Here, we describe a new mechanistic link between mitochondria and cardiac morphogenesis, uncovered by studying mice with systemic loss of the mitochondrial citrate carrier SLC25A1. *Slc25a1* null embryos displayed impaired growth, cardiac malformations, and aberrant mitochondrial function. Importantly, *Slc25a1* heterozygous embryos, which are overtly indistinguishable from wild type, exhibited an increased frequency of these defects, suggesting *Slc25a1* haploinsuffiency and dose-dependent effects. Supporting clinical relevance, we found a near-significant association between ultrarare human pathogenic *SLC25A1* variants and pediatric congenital heart disease. Mechanistically, SLC25A1 may link mitochondria to transcriptional regulation of metabolism through epigenetic control of gene expression to promote metabolic remodeling in the developing heart. Collectively, this work positions SLC25A1 as a novel mitochondrial regulator of ventricular morphogenesis and cardiac metabolic maturation and suggests a role in congenital heart disease.

## INTRODUCTION

In the postnatal heart, mitochondria drive metabolism by providing oxidative energy to fuel contraction. However, the prenatal heart is a hypoxic environment ^1^ with less clear roles for mitochondrial oxidative energy systems. Nonetheless, mitochondrial metabolism is critical for embryogenesis, as systemic defects in respiratory chain regulation result in embryonic lethality by embryonic (E) day 10.5 (E10.5) ^2–6^. The early embryonic heart (before E11.5) exhibits structurally and functionally immature mitochondria and relies mainly on glycolysis for ATP. As development proceeds (E14.5), cardiac mitochondria mature, and embryonic cardiac metabolism shifts toward mitochondrial oxidation ^7, 8^. This metabolic transition may support the increasing energetic demands of the developing heart since increased oxidative metabolism is a defining feature of cardiomyocyte maturation ^9^.

In parallel with this embryonic metabolic shift, the developing heart undergoes key processes of ventricular wall maturation including trabeculation, septation, and compaction ^10, 11^. While metabolism and ventricular morphogenesis are seemingly disparate processes, human primary mitochondrial disorders may offer insight to an intriguing connection between mitochondrial energetics and ventricular morphogenesis. Patients with mitochondrial disorders including Barth syndrome, MELAS (mitochondrial encephalopathy, lactic acidosis, and stroke-like episodes), and MERRF (myoclonic epilepsy with ragged-red fibers) can present with left ventricular noncompaction, a congenital heart defect caused by impaired compaction of the ventricular wall, suggesting an association between oxidative phosphorylation defects and cardiac structural malformations ^12, 13^. At present, the molecular mechanisms connecting mitochondrial energetics to the tightly regulated process of ventricular wall morphogenesis are not fully understood. However, defining these connections may help inform the basic biology of cardiac development and highlight new genetic targets for improved screening or interventions for congenital heart disease (CHD).

One possible connection between mitochondrial energetics and other cellular processes is the mitochondrial citrate carrier SLC25A1. SLC25A1 is a mitochondrial inner membrane transporter that exports citrate out of the mitochondrial matrix ^14, 15^. While citrate in the mitochondria is a key intermediate of the tricarboxylic acid (TCA) cycle and thus central to mitochondrial oxidative metabolism, citrate in the cytosol is converted into acetyl-CoA, a substrate that can be used in various biological pathways ranging from lipid biosynthesis to protein post-translational acetylation and epigenetic control of gene expression ^16–18^.

Loss of SLC25A1 reduces respiratory chain function in vitro ^19–21^, suggesting that SLC25A1 also may directly regulate mitochondrial function. The *SLC25A1* gene is located within a region of chromosome 22 that is compromised in 22q11.2 deletion syndrome (22q11.2DS, also known as DiGeorge syndrome). Individuals missing this segment on one copy of chromosome 22 exhibit a multisyndromic disorder that includes congenital heart defects in approximately 60-80% of affected individuals ^22, 23^. Interestingly, we reported that the neurodevelopmental pathology of 22q11.2DS arises in part due to loss of SLC25A1, which leads to mitochondrial dysfunction and the dysregulation of an integrated network of mitochondrial proteins ^24^. Collectively, these data indicate SLC25A1 is a mitochondrial protein that can both regulate mitochondrial function and connect mitochondria to cytosolic processes. However, the roles of SLC25A1 in vivo, particularly during development, are not fully delineated.

Here we investigated the role of SLC25A1 in development using a systemic *Slc25a1* knockout (KO) mouse model. Our findings implicate SLC25A1 in essential roles in epigenetic regulation of metabolic reprogramming in the developing heart and cardiac morphogenesis. We further identify a near-significant association of ultrarare human *SLC25A1* variants with pediatric CHD. Importantly, this work suggests that SLC25A1 is a novel mitochondrial regulator of cardiac morphogenesis and that the *Slc25a1* KO mouse model recapitulates the cardiac defects of 22q11.2DS, providing important avenues to new insight for this condition.

## RESULTS

### Systemic *Slc25a1* deletion in mice impairs embryonic growth and causes perinatal lethality

We used the *Slc25a1* KO mouse line developed by the International Mouse Phenotyping Consortium to model systemic deletion of *Slc25a1* ^25^, and importantly, to reflect systemic gene loss that would be observed in humans with SLC25A1 deficiencies. For these mice, the *Slc25a1* KO allele (*Slc25a1*^−^) was generated by insertion of an IRES:LacZ trapping cassette and a neomycin selection cassette between exons 1 and 2 of the *Slc25a1* locus (Fig. 1a). *Slc25a1* systemic KO mice (*Slc25a1^−/−^*) exhibit perinatal lethality ^26^. Thus, we conducted timed matings of *Slc25a1* hemizygous KO mice (*Slc25a1^+/−^*) to study the embryonic effects of complete *Slc25a1* ablation (*Slc25a1^−/−^*) and *Slc25a1* hemizygous loss (*Slc25a1^+/−^*), compared with wild type (WT) littermate controls (*Slc25a1^+/+^*). We confirmed that *Slc25a1^−/−^* embryos at E18.5 were viable, and *Slc25a1^+/−^* intercrosses yielded the anticipated genotypic frequencies at this stage (Fig. 1f). However, E18.5 *Slc25a1^−/−^* embryos displayed impaired growth with significantly decreased crown-rump length compared to *Slc25a1^+/−^* and *Slc25a1^+/+^* littermate controls (Fig. 1b,c). Additionally, E18.5 *Slc25a1^−/−^* embryos displayed microencephaly, anencephaly, encephalocele, or gastroschisis (data not shown). Immunoblot analyses of SLC25A1 protein levels in E18.5 heart and brain tissues revealed gene dosage-dependent expression: SLC25A1 expression was robust in *Slc25a1^+/+^* embryos, reduced in *Slc25a1^+/−^* embryos, and absent from *Slc25a1^−/−^* embryos (Fig. 1d). Despite the severe effects of full *Slc25a1* deletion, E18.5 *Slc25a1^+/−^* embryos were morphologically indistinguishable from *Slc25a1^+/+^* controls.

**Figure 1.**
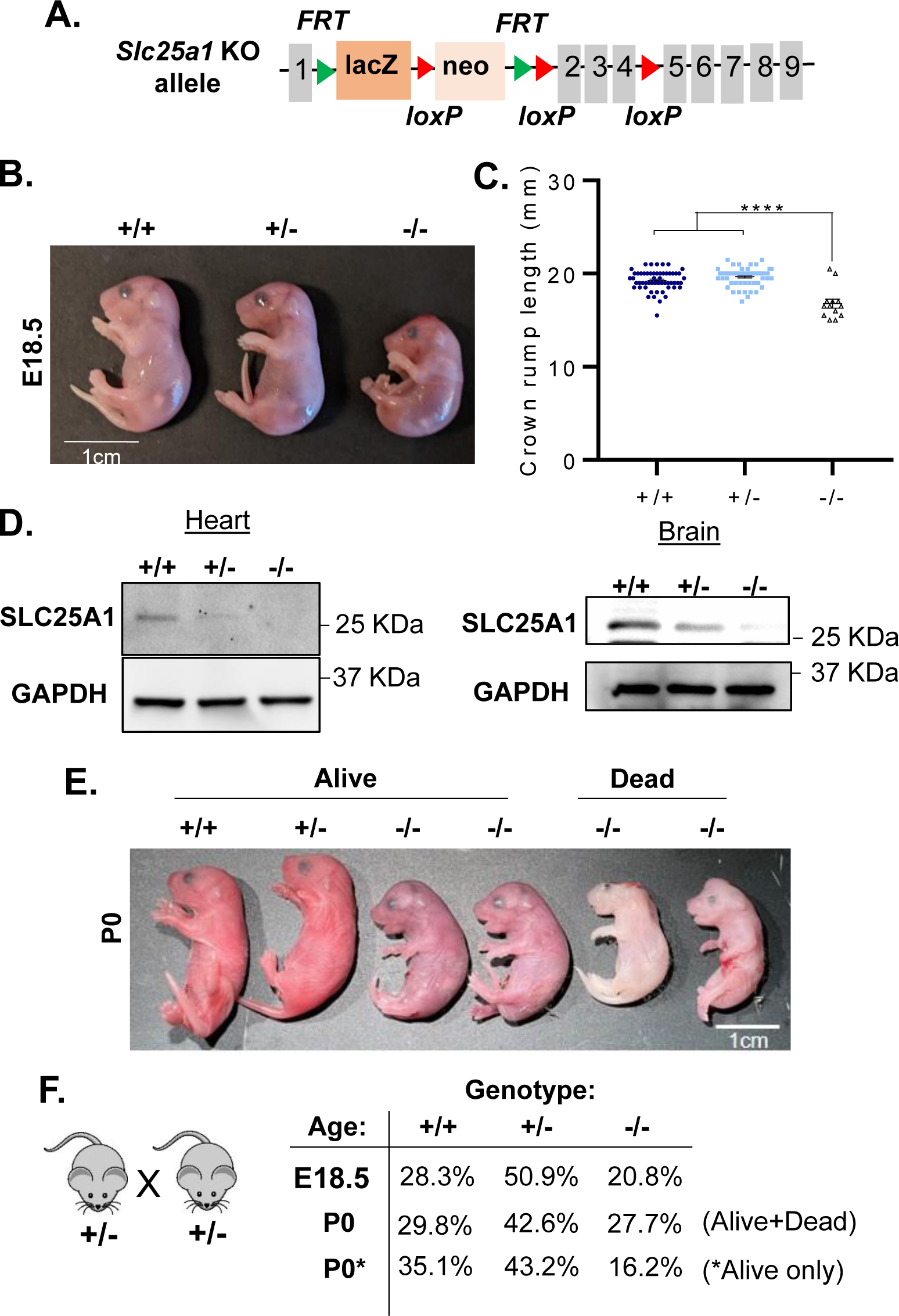
Systemic *Slc25a1* deletion impairs mouse embryonic growth and causes perinatal lethality. (A) Schematic of the *Slc25a1* knockout (KO) first allele. A lacZ-neo KO cassette was inserted between exons 1 and 2 of *Slc25a1*. (B) Representative E18.5 *Slc25a1^+/+^*, *Slc25a1^+/−^*, and *Slc25a1^−/−^* embryos obtained from timed intercross of *Slc25a1^+/−^* mice. (C) Quantification of crown-rump lengths of E18.5 embryos of the indicated genotypes (n = 13– 26/group). Values reported as mean ± SEM. Student’s *t* test was used for statistical analysis. ****P < 0.0001. (D) Western blot of SLC25A1 expression in total protein isolated from E17.5 hearts and brains from the indicated genotypes. GAPDH is a loading control. (E) Representative image of P0 pups obtained from timed intercross of *Slc25a1^+/−^* mice. (F) Genotype frequencies of E18.5 embryos and P0 pups obtained from timed intercross of *Slc25a1^+/−^* mice.

In line with previous reports, we were unable to recover adult *Slc25a1^−/−^* mice from *Slc25a1^+/−^* intercrosses. However, *Slc25a1^−/−^* pups were born and obtained at the anticipated genotype frequencies (Fig. 1e,f). Further, postnatal (P) day 0 (P0) *Slc25a1^−/−^* neonates were small and cyanotic, and some displayed microencephaly and gastroschisis (Figure 1e). Importantly, *Slc25a1^−/−^* neonates did not survive beyond P0 and died within 6 hours of birth, suggesting that systemic ablation of *Slc25a1* causes perinatal lethality.

### Partial *Slc25a1* deletion is sufficient to produce cardiac structural defects in mice

SLC25A1 is highly expressed in the mouse heart ^27, 28^, and given that P0 *Slc25a1^−/−^* neonates were cyanotic, we hypothesized that SLC25A1 loss could impair cardiac contractile function. We assessed SLC25A1 expression in the developing heart by immunofluorescence in WT E10.5, E12.5, and E14.5 embryos and E18.5 hearts. Because the three layers of the cardiac wall (endocardium, myocardium, and epicardium) play distinct roles in ventricular morphogenesis, we investigated expression of SLC25A1 in the different layers of the heart. Examining fields of the embryonic heart spanning the myocardium and epicardium (Fig. 2a) as well as the myocardium and endocardium (Supplemental Figure 1), we observed strong SLC25A1 expression in the myocardium and epicardium (Fig. 2a) and minimal expression in the endocardium (Supplemental Figure 1).

**Figure 2.**
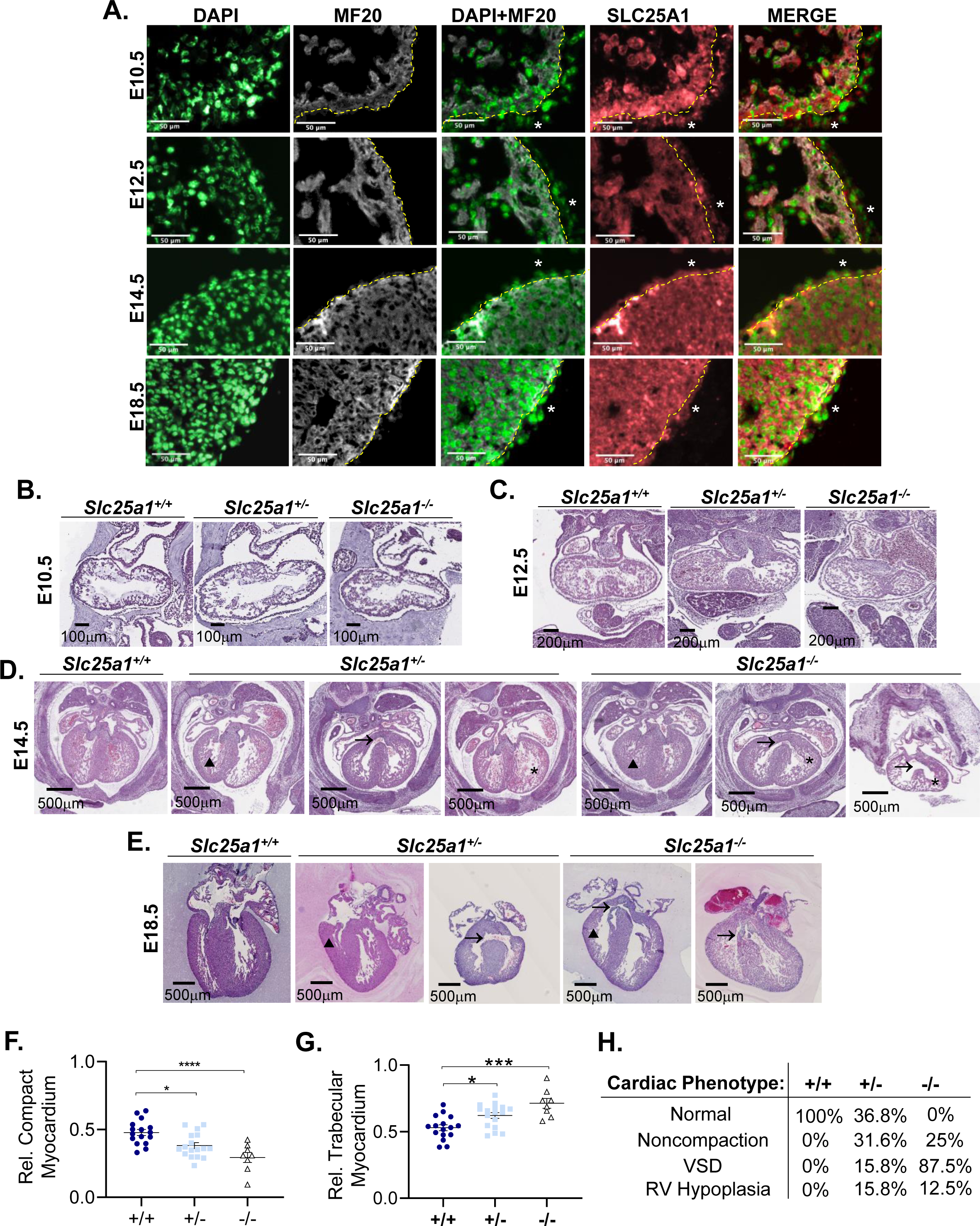
Partial *Slc25a1* deletion is sufficient to produce cardiac structural defects in mice. (A) Representative immunofluorescence images of SLC25A1 expression in wild type embryo (E10.5, E12.5, and E14.5) and heart (E18.5) sections (6 μm). DAPI was used to counterstain nuclei, and MF20 was used to counterstain cardiomyocytes. Dashed yellow line outlines myocardium from the epicardium; asterisk indicates epicardium positive for SLC25A1. (B) Representative images of E10.5 embryos, (C) E12.5 embryos, (D) E14.5 embryos, and (E) E18.5 hearts from the indicated genotypes stained with hematoxylin and eosin. Asterisk indicates noncompaction, arrow indicates ventricular septal defect, and triangle indicates right ventricular hypoplasia. (F) Quantification of relative compact myocardium thickness or (G) relative trabecular myocardium thickness in E18.5 hearts (n=8-17/group). (H) Frequency of cardiac defects observed at E14.5 in embryos with the indicated genotypes (n = 27/group). Values reported as mean ± SEM. Student’s *t* test was used for statistical analysis. *P < 0.05; ***P < 0.0005; ****P < 0.0001.

To investigate the structural consequences of SLC25A1 loss in the embryonic heart, we conducted timed matings of *Slc25a1^+/−^* mice to obtain *Slc25a1^−/−^*, *Slc25a1^+/−^*, and control *Slc25a1^+/+^* embryos. Embryos were collected at E10.5, E12.5, and E14.5 and embryonic hearts were collected at E18.5 to reflect timepoints during ventricular morphogenesis when trabeculation occurs (E10.5 and E12.5), when septation should be complete and ventricular wall compaction should be fulminant (E14.5), and when the major embryonic milestones of cardiac morphogenesis should be complete (E18.5) ^10, 11^. Histological analyses of serial embryo sections showed grossly similar cardiac morphology of E10.5 *Slc25a1^+/+^, Slc25a1^+/−^, and Slc25a1^−/−^* embryos (Fig. 2b). At E12.5, the compact myocardium of *Slc25a1^−/−^* embryos appeared to be slightly less developed than that of *Slc25a1^+/+^* and *Slc25a1^+/−^* embryos (Fig. 2c). By E14.5 and E18.5, however, hearts from *Slc25a1^−/−^* embryos displayed striking cardiac abnormalities (Fig. 2d–h) including ventricular septal defects (VSDs); ventricular noncompaction, as evidenced by decreased relative compact myocardium thickness (Fig. 2f) and increased relative trabecular myocardium thickness (Fig. 2g) compared to age-matched *Slc25a1^+/+^* controls; and right ventricular hypoplasia. As VSDs have been reported in *Slc25a1* KO embryos ^29^, our findings further extend the cardiac abnormalities associated with *Slc25a1* deletion to include noncompaction and right ventricular hypoplasia.

*Slc25a1^+/−^* embryos, which were grossly indistinguishable from *Slc25a1^+/+^* littermate controls, also unexpectedly displayed increased frequency of cardiac malformations including VSDs, ventricular noncompaction, and right ventricular hypoplasia (Fig. 2d–h). Collectively, our data demonstrate that SLC25A1 is necessary for ventricular morphogenesis and that heterozygous loss (thus *Slc25a1* haploinsufficiency) is sufficient to cause structural defects in the developing heart.

### *Slc25a1* haploinsufficiency decreases neonatal survival in mice

While *Slc25a1^+/−^* embryos displayed congenital heart defects, adult *Slc25a1^+/−^* mice were viable and fertile and did not display differences in survival with aging (data not shown). This disconnect prompted us to further investigate the structural and functional consequences of *Slc25a1* haploinsufficiency in the postnatal adult heart, specifically if adult *Slc25a1^+/−^* mice also harbor cardiac defects. Histological analyses of hearts from 2-month-old *Slc25a1^+/−^* and *Slc25a1^+/+^* control adult mice (n = 10/group) revealed no structural or morphological defects (Supplemental Figure 2A). Additionally, no changes were observed in gravimetrics (heart weight to body weight ratios; Supplemental Figure 2B) or echocardiographic parameters (Supplemental Figure 2C), suggesting that cardiac structure and function were unaltered in adult mice with *Slc25a1* haploinsufficiency.

To understand how we could observe cardiac malformations in *Slc25a1^+/−^* embryos but not adult mice, we examined the frequencies of progeny genotypes. For colony maintenance, we used a breeding strategy of *Slc25a1^+/−^* mice crossed to *Slc25a1^+/+^* WT mice, with genotyping conducted on tissue biopsies obtained at P7. Genotype analyses of 446 pups from 80 litters revealed a slight but significant [χ^2^ (1, N = 446) = 6.0628; p = 0.014] increase in the frequency of P7 *Slc25a1^+/+^* (56% observed; 50% expected) versus *Slc25a1^+/−^* pups obtained (44% observed; 50% expected; Supplemental Figure 2D), even though *Slc25a1^+/−^* pups were recovered at expected frequencies at P0 (N=47, Figure 1F). Thus, our data suggest that *Slc25a1* haploinsufficiency decreases neonatal survival. Moreover, as cardiac morphometry and function in *Slc25a1^+/−^* mice were normal at 2 months of age, our data suggest that *Slc25a1^+/−^* pups born with cardiac defects die in the neonatal period.

### Association of ultrarare *SLC25A1* variants with pediatric CHD

In humans, *SLC25A1* single-gene mutations underlie two rare neurometabolic and neuromuscular diseases, D-2 and L-2 aciduria (DL-2HGA; OMIM: 615182) and congenital myasthenic syndrome-23 (CMS23; OMIM: 618197) ^30–32^. There are no human cardiac phenotypes annotated to *SLC25A1* defects (https://hpo.jax.org/app/browse/gene/6576), and it is unknown if *SLC25A1* mutations underlie cardiac disease. Our surprising findings of ventricular malformations in *Slc25a1^+/−^* and *Slc25a1^−/−^* mouse embryos prompted us to investigate if *SLC25A1* mutations are associated with CHD in humans. We analyzed published whole-exome sequencing data from the Pediatric Cardiac Genomics Consortium (PCGC), encompassing 3740 CHD probands with 3401 pseudo-sibship case-controlled pairs (generated from parent-proband trios) ^33, 34^. Rare *SLC25A1* variants were selected using a minor allele frequency of <0.001 in the Genome Aggregation Database (gnomAD) and a combined annotation-dependent depletion score of >20. A separate set including only ultrarare *SLC25A1* variants (that do not appear in gnomAD exomes) was generated. Interestingly, while we were unable to identify an association of rare and ultrarare *SLC25A1* variants with specific CHD presentations (data not shown), gene burden analyses using the kernel-based adaptive cluster method revealed that ultrarare nonsynonymous and stop-gain *SLC25A1* variants were associated with CHD at just above the significance threshold (p = 0.058; Supplemental Table 1). These data substantiate our mouse studies and provide clinical relevance for the role of SLC25A1 in the developing heart.

### *Slc25a1* deletion alters metabolic gene expression in mice

SLC25A1 has a well-established function as a mitochondrial solute transporter that exports mitochondrial citrate ^14, 15^. Work in cancer cells as well as neuronal systems suggests a role for SLC25A1 beyond metabolite transport in direct regulation of respiratory chain function, as SLC25A1 deletion in these systems is associated with impaired respiration ^19, 24, 35^. Our work in the HAP1 human near-haploid lymphoblastoid cell line offers a potential mechanism connecting SLC25A1 to respiratory chain function through regulation of mitochondrial translation, as loss of SLC25A1 is associated with reduced expression of core mitochondrial ribosome subunits ^21^. Thus, we hypothesized that SLC25A1-dependent regulation of mitochondrial translation and mitochondrial dysfunction would also be operant in the developing heart. We examined protein expression levels of key mitochondrial ribosome subunits, MRPS22 and MRPS18B, in E17.5 *Slc25a1^+/+^*, *Slc25a1^+/−^*, and *Slc25a1^−/−^* embryonic hearts. Surprisingly, MRPS22 and MRPS18B levels were unchanged with *Slc25a1* deletion (Supplemental Figure 3), suggesting that SLC25A1 does not regulate mitochondrial translation in the developing heart.

To delineate the overall consequences of *Slc25a1* deletion on metabolism, we used a NanoString mRNA quantification panel to profile expression of 768 metabolism-related genes encompassing key metabolic pathways. Profiling metabolic gene expression in *Slc25a1^+/−^* versus *Slc25a1^+/+^, Slc25a1^−/−^* versus *Slc25a1^+/+^*, *and Slc25a1^+/−^* versus *Slc25a1^−/−^* hearts revealed a discrete set of 28 differentially expressed genes with *Slc25a1* deletion at E17.5 (p < 0.05). Of these genes, 10 were differentially expressed between *Slc25a1^−/−^* versus *Slc25a1^+/+^*, two between *Slc25a1^+/−^* versus *Slc25a1^+/+^*, and 11 between *Slc25a1^+/−^* versus *Slc25a1^−/−^* hearts (fold-change = 1.5; p < 0.05; Fig. 3a). We used these differentially expressed genes to identify ontologies with the Metascape tool ^36^ and identified antigen processing, organic acid transport, positive regulation of cell death, AMPK signaling, and response to hormone as the most significantly altered pathways with *Slc25a1* deletion (Fig. 3b).

**Figure 3.**
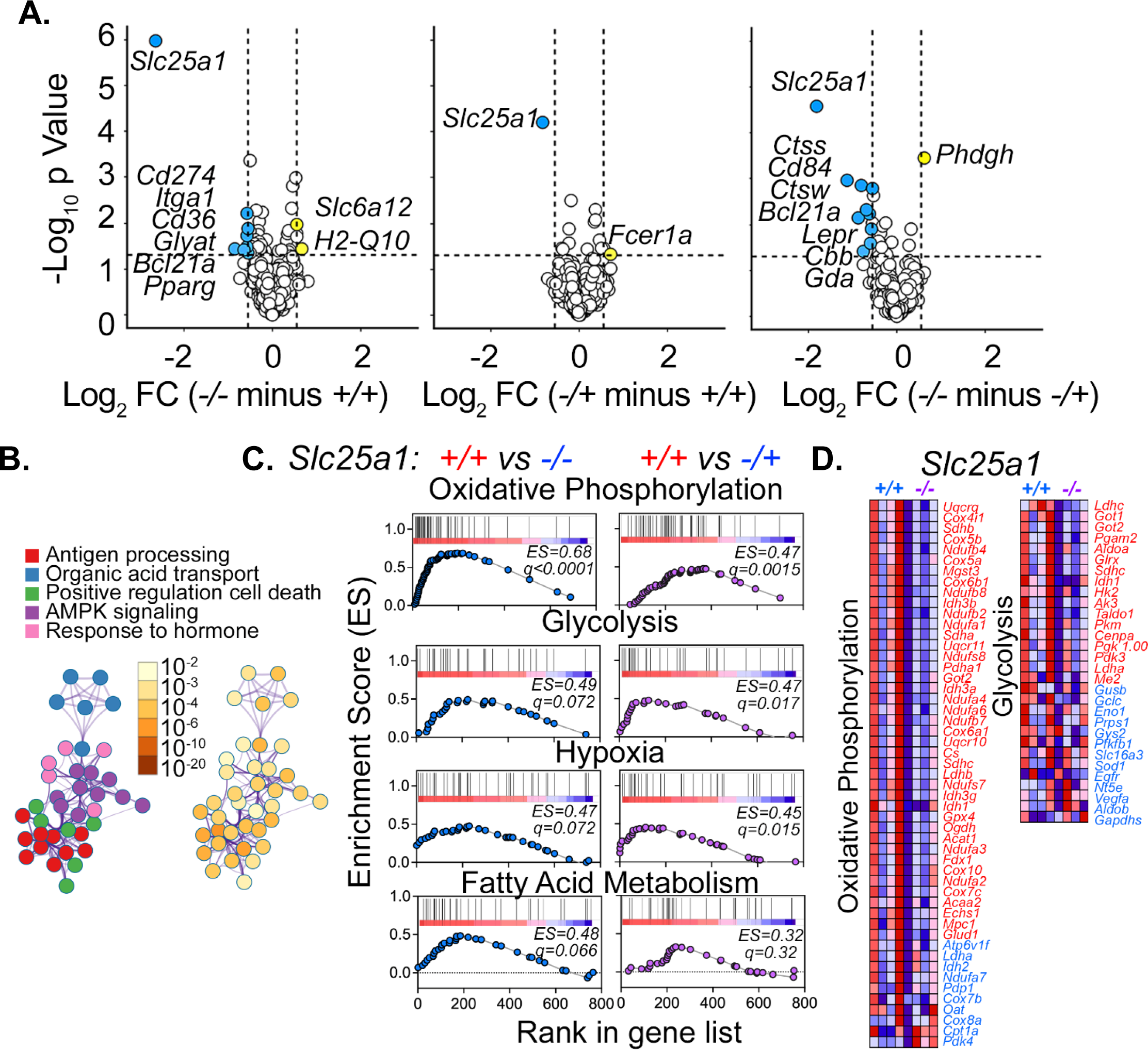
*Slc25a1* deletion alters metabolic gene expression in mice. (A) Volcano plots of Nanostring metabolic transcript profiling comparing gene expression of *Slc25a1^−/−^* and *Slc25a1^+/+^*, *Slc25a1^+/−^* and *Slc25a1^+/+^,* and *Slc25a1^−/−^* and Slc25a1^+/−^ E17.5 hearts. Blue denotes downregulated transcripts. Yellow denotes upregulated transcripts. (B) Gene ontologies identified with Metascape from genes differentially expressed in *Slc25a1^−/−^* and *Slc25a1^+/−^* versus *Slc25a1^−/−^* hearts. (C) Geneset enrichment analysis of oxidative phosphorylation, glycolysis, hypoxia, and fatty acid metabolism genes in *Slc25a1^+/+^* vs. *Slc25a1^−/−^* and *Slc25a1^+/+^* vs. *Slc25a1^+/−^* E17.5 hearts. (D) Heatmaps visualizing gene expression of oxidative phosphorylation and glycolysis genes from Nanostring transcript profiling in E17.5 *Slc25a1^+/+^* vs. *Slc25a1^−/−^* hearts. Blue denotes reduced expression; red denotes increased expression.

To further define the metabolic processes impacted by *Slc25a1* deletion, we performed gene set enrichment analysis (GSEA) of REACTOME pathways using normalized gene expression from the NanoString panel as gene ranks. GSEA revealed oxidative phosphorylation, glycolysis, hypoxia, and fatty acid metabolism as the most significantly altered pathways in *Slc25a1^−/−^* versus *Slc25a1^+/+^* and *Slc25a1^+/−^* versus *Slc25a1^+/+^* hearts, with higher enrichment scores for these pathways in control *Slc25a1^+/+^* hearts versus hearts with *Slc25a1* deletion (*Slc25a1^−/−^* and *Slc25a1^+/−^*) (Fig. 3c,d). These results suggest that *Slc25a1* deletion induces mitochondrial dysfunction and metabolic remodeling of the embryonic heart.

More importantly, these data also show that loss of only one copy of *Slc25a1* is sufficient to induce gene expression changes. Intriguingly, comparisons of the enrichment plots for oxidative phosphorylation, glycolysis, hypoxia, and fatty acid metabolism revealed a right-ward shift of the *Slc25a1^+/−^* versus *Slc25a1^+/+^* plots as compared to *Slc25a1^−/−^* versus *Slc25a1^+/+^* plots (Fig. 3c). These graded differences between homozygous and heterozygous *Slc25a1* knockouts suggest a copy number-dependent degree of severity in mitochondrial and metabolic dysregulation, such that full *Slc25a1* ablation induces greater dysregulation than partial loss. These results align with our findings that *Slc25a1^−/−^* embryos displayed a higher frequency and severity of ventricular malformations than *Slc25a1^+/−^* embryos.

### *Slc25a1* deletion causes mitochondrial ultrastructural defects in the developing mouse heart

Our metabolic gene expression profiling suggested that SLC25A1 loss causes mitochondrial dysfunction in the developing heart in a *Slc25a1* dosage-dependent manner. Because mitochondrial structure and function are often linked ^37^, we investigated the impact of SLC25A1 loss on mitochondrial ultrastructure. Transmission electron microscopy on *Slc25a1^−/−^*, *Slc25a1^+/−^*, and *Slc25a1^+/+^* hearts collected at E17.5 revealed both partial and complete *Slc25a1* ablation induced striking alterations in cardiac mitochondrial morphology—mitochondria from *Slc25a1^+/−^* and *Slc25a1^−/−^* hearts displayed highly immature and disordered cristae as well as open matrix voids (Fig. 4a). Quantification of mitochondrial morphology further revealed that *Slc25a1^+/−^* and *Slc25a1^−/−^* mitochondria had significantly increased area (Supplemental Figure 4A) and minimal Feret’s diameter (Supplemental Figure 4B), despite similar circularity (Supplemental Figure 4C), suggesting larger and swollen mitochondria. Mitochondria from *Slc25a1^−/−^* hearts further displayed significantly decreased aspect ratio (Supplemental Figure 4D) and increased roundness (Supplemental Figure 4E), suggesting greater changes in mitochondrial morphology, compared to *Slc25a1^+/−^* and *Slc25a1^+/+^* hearts. Our data suggest that, similar to metabolic gene expression changes, mitochondrial ultrastructural abnormalities are *Slc25a1* dosage-dependent. Moreover, as cardiac mitochondria become more elongated, develop more elaborate and organized cristae, and become more structurally complex during cardiac development, our data suggest that SLC25A1 loss impairs mitochondrial structural maturation.

**Figure 4.**
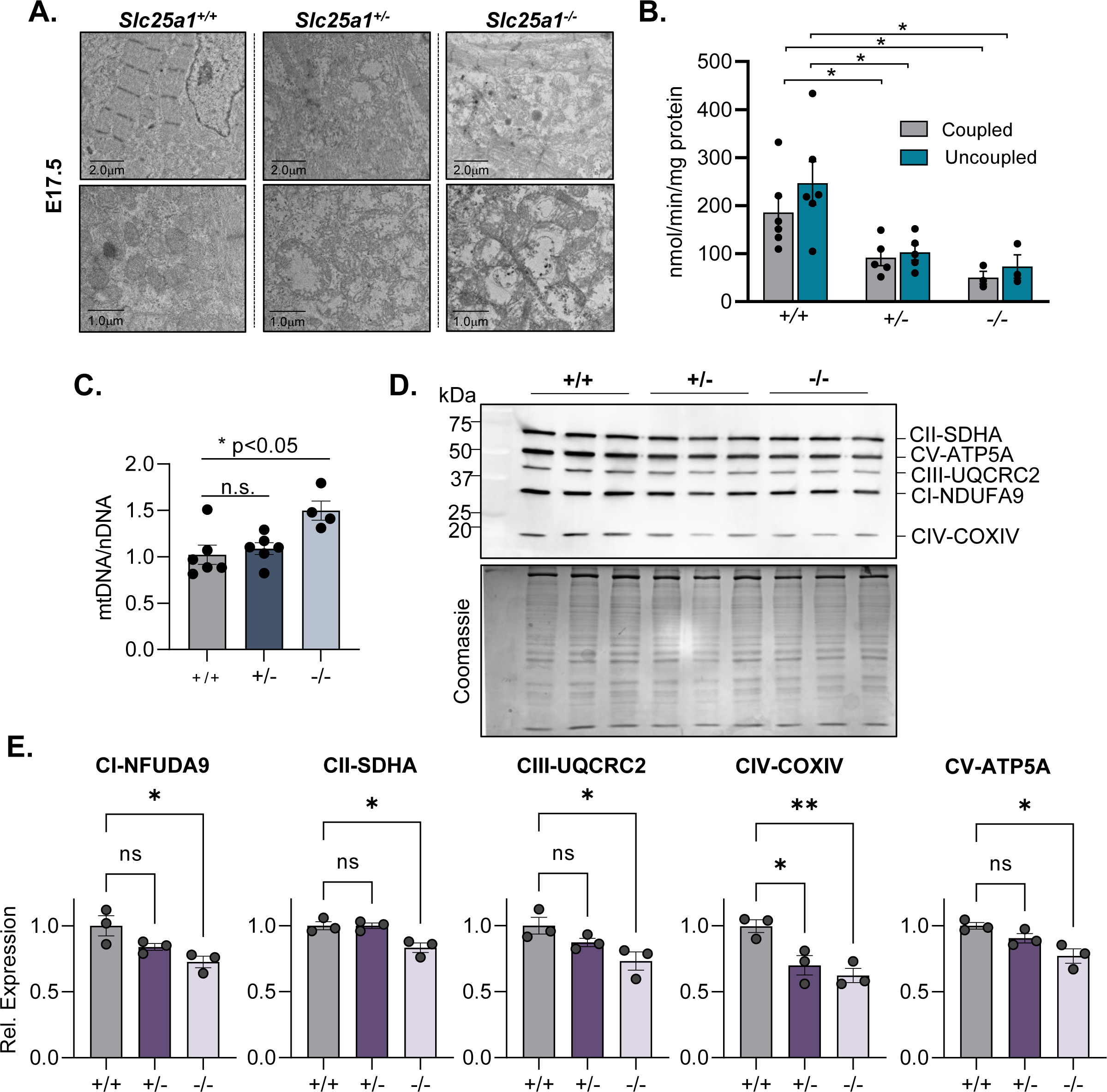
*Slc25a1* deletion impairs respiratory chain function in the developing mouse heart. (A) Representative electron micrographs of cardiac ultrastructure from the indicated genotypes at E17.5 (n = 3 *Slc25a1^+/+^*; n = 7 *Slc25a1^+/−^*; n = 3 *Slc25a1^−/−^* embryos analyzed). (B) Coupled and uncoupled mitochondrial oxygen consumption rates measured in lightly permeabilized whole E17.5 hearts of the indicated genotypes (n = 5 *Slc25a1^+/+^*; n = 5 *Slc25a1^+/−^*; n = 3 *Slc25a1^−/−^* hearts analyzed). (C) Quantification of mitochondrial DNA (mtDNA) to nuclear DNA (nDNA) ratios in E17.5 hearts of the genotypes indicated (n = 6 *Slc25a1^+/+^*; n = 6 *Slc25a1^+/−^*; n = 4 *Slc25a1^−/−^* embryos analyzed). (D) Western blot of respiratory chain complex subunits (SDHA, ATP5A, UQCRC2, NDUFA9, and COXIV). Coomassie staining of the gel was used as a protein loading control. (E) Densitometry quantification of NDUFA9, SDHA, UQCRC2, COXIV, and ATP5A expression (n = 3/group). Values presented as means ± SEM. Student’s *t* test was used for statistical analysis. ns denotes not significant; *P < 0.05; **P < 0.005.

### *Slc25a1* deletion impairs respiratory chain function in the developing mouse heart

As our data suggested that *Slc25a1* deletion causes mitochondrial dysfunction, we assessed the impact of *Slc25a1* deletion on mitochondrial respiratory chain function. Respirometry was conducted on hearts isolated from E17.5 *Slc25a1^+/−^*, *Slc25a1^−/−^*, and *Slc25a1^+/+^* embryos. Consistent with our gene expression data, coupled mitochondrial oxygen consumption (supported by glutamate and malate) of *Slc25a1^+/−^* and *Slc25a1^−/−^* hearts was significantly decreased compared to *Slc25a1^+/+^* controls (Figure 4B). These respiration deficits were also observed in the presence of chemical uncoupling by FCCP (carbonyl cyanide-p-trifluoromethoxyphenylhydrazone), indicating that *Slc25a1* deletion also impairs maximal respiratory capacity. Together, these results support mitochondrial dysfunction and impaired oxidative phosphorylation with SLC25A1 loss.

Because defects in mitochondrial cristae ultrastructure have been associated with mitochondrial DNA (mtDNA) depletion ^38, 39^ and the mtDNA encodes essential respiratory chain complex subunits, *Slc25a1* deletion-induced mitochondrial ultrastructural abnormalities (Figure 4A) could lead to respiration defects by causing mtDNA loss. Thus, we measured mtDNA to nuclear DNA (nDNA) ratios (mtDNA/nDNA) in total DNA isolated from E17.5 *Slc25a1^+/−^*, *Slc25a1^−/−^*, and *Slc25a1^+/+^* hearts. While mtDNA/nDNA ratios were not significantly different between *Slc25a1^+/−^* and *Slc25a1^+/+^* control hearts, mtDNA/nDNA ratios were elevated in *Slc25a1^−/−^* hearts, indicating increased mtDNA content (Fig. 4c). Thus, SLC25A1 deletion-associated respiration defects are not due to mtDNA depletion.

To further confirm that SLC25A1 deletion does not alter mitochondrial content in the developing heart, we conducted immunoblotting for TOM20—a ubiquitously expressed subunit of the mitochondrial import machinery that is often used as a mitochondrial housekeeping gene—and observed no differences between *Slc25a1^+/−^*, *Slc25a1^−/−^*, and *Slc25a1^+/+^* control hearts (Supplemental Figures 5A,B). Collectively, these data suggest that SLC25A1 expression does not impact overall cardiac mitochondrial content.

We also directly investigated the impact of *Slc25a1* deletion on respiratory chain complex subunit expression (NDUSFA9 of complex I, SDHA of complex II, UQCRC2 of complex III, COXIV of complex IV, and ATP5A of complex V) by immunoblotting total heart lysates from E17.5 *Slc25a1^+/−^*, *Slc25a1^−/−^*, and *Slc25a1^+/+^* embryos (Fig. 4d). *Slc25a1^+/−^* hearts displayed a trend for decreased NDUSFA9, UQCRC2, and ATP5A levels as well as significantly decreased COXIV levels compared to controls (Fig. 4e). *Slc25a1^−/−^* hearts displayed significantly decreased expression of all respiratory chain subunits compared to controls (Fig. 4e). Importantly, the *Slc25a1* dosage-dependent decrease in respiratory chain subunit expression was consistent with dosage-dependent alterations in metabolic gene expression, mitochondrial ultrastructural abnormalities and impaired respiration in SLC25A1-deleted hearts.

### Computational modeling predicts metabolic dysregulation and increased glucose dependence with *Slc25a1* deletion

To investigate the overall metabolic consequences of *Slc25a1* deletion in the E17.5 mouse heart, we performed targeted gas chromatography–mass spectrometry (GC-MS) metabolomic profiling on 25 key metabolites, encompassing TCA cycle intermediates, amino acids, and metabolic byproducts like lactate and 2-hydroxyglutaric acid, that are critical to cardiac energetics (Figure 5A). Principal component analysis (PCA) as well as unbiased hierarchal clustering of the measured analytes separated embryonic heart samples according to their *Slc25a1* genotype (Fig 5b,c), with *Slc25a1^+/−^* hearts displaying an intermediate metabolite profile compared to *Slc25a1^+/+^* and *Slc25a1^−/−^* hearts. Notably, citrate was significantly less abundant in *Slc25a1^−/−^* compared to *Slc25a1^+/+^* hearts, while *Slc25a1^+/−^* hearts displayed a trend toward reduced citrate (Fig. 5d), indicating that SLC25A1 deletion causes an overall decrease in cardiac citrate levels. Additionally, glutamate, cystine and lysine abundance showed a pronounced increase with SLC25A1 deletion (Fig. 5d), while threonine, tryptophan, aconitate and pyroglutamate levels were decreased. Glutamine and glutamate undergo cyclization to pyroglutamate during the de novo synthesis of gamma-glutamyl cycle of glutathione (l-*γ*-glutamyl-l-cysteinyl-glycine; GSH). Likewise, cystine is a precursor for the synthesis of GSH. Hence, the marked reduction in pyroglutamate and increased abundance of glutamate, as well as cystine indicates reduced GSH synthesis in response to *Slc25a1* deletion.

**Figure 5.**
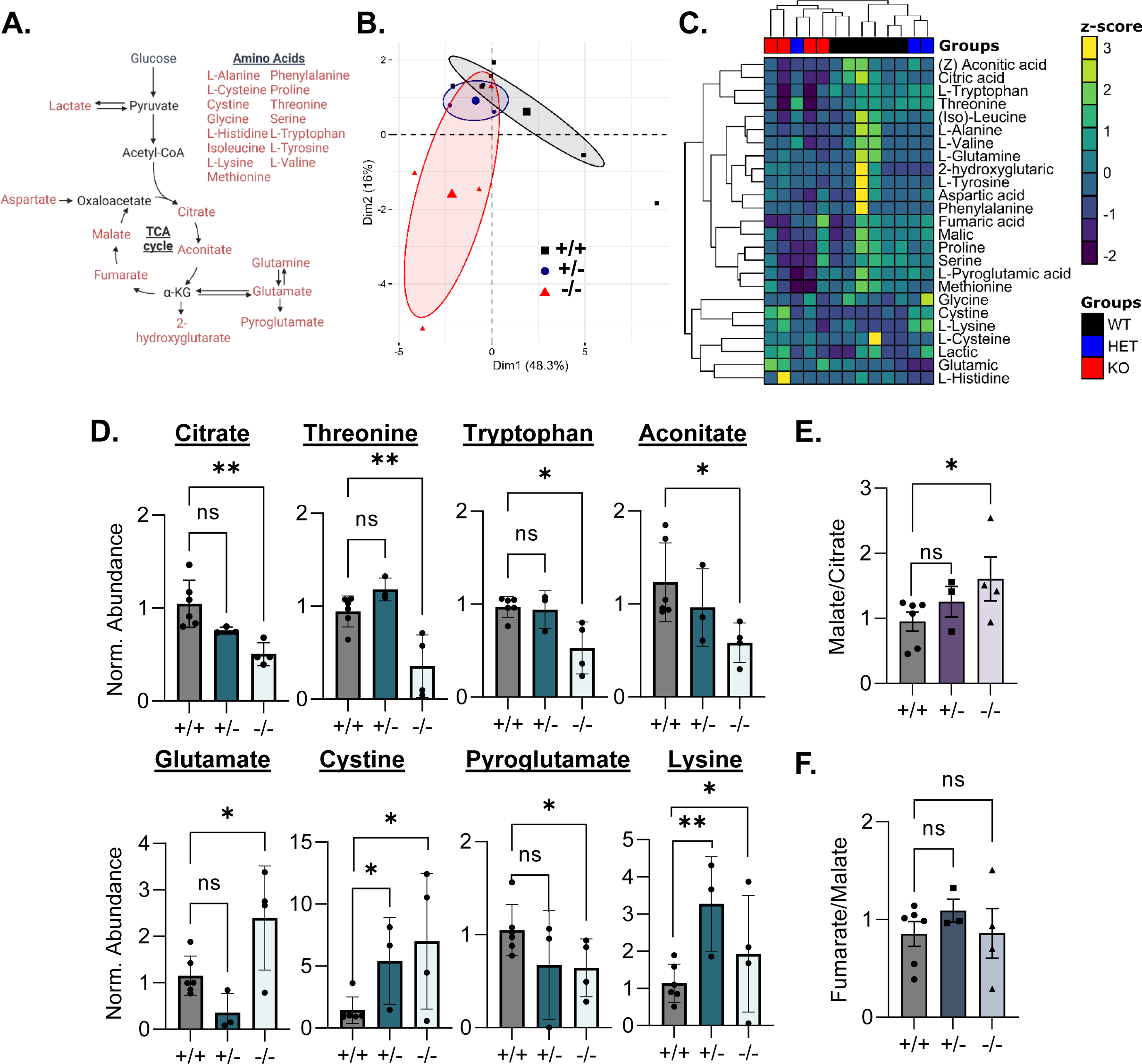
Targeted metabolomic profiling reveals amino acid dysregulation and impaired Krebs cycle function with SLC25A1 deletion. (A) Schematic of the metabolites profiled by GC-MS. (B) Principal component analysis (PCA) and (C) unbiased hierarchical clustering of profiled analytes showing clear segregation of *Slc25a1^+/+^*(n=6), *Slc25a1^+/−^* (n=3), and *Slc25a1^−/−^* (n=4) hearts. (D) Normalized abundance of selected metabolites that are significantly altered in HET and KO hearts. Values presented as means ± SEM. Student’s *t* test was used for statistical analysis. P < 0.05 was considered significant, *P < 0.05 and **P<0.005. (E) Malate-to-citrate ratios and (F) Fumarate-to-malate ratios in *Slc25a1^+/+^*, *Slc25a1^+/−^* and *Slc25a1^−/−^* hearts. Values presented as means ± SEM. One-way ANOVA followed by a post hoc Dunnett’s test was used for statistical analysis. P < 0.05 was considered significant, ns denotes not significant, *P < 0.05.

Because glucose-derived citrate can either be exported from the mitochondria into the cytosol via SLC25A1 or further decarboxylated to succinate followed by oxidation to malate as part of the TCA cycle, we compared the malate-to-citrate and fumarate-to-malate ratios to assess the incorporation and conversion of citrate-derived carbons at different levels within the TCA cycle. With *Slc25a1* deletion, the malate-to-citrate ratio was significantly increased in *Slc25a1^−/−^* hearts compared to controls, while *Slc25a1^+/−^* hearts displayed a trend for an increase (Fig. 5e), while the fumarate-to-malate ratios remained unchanged (Fig. 5f). Together these data suggest that SLC25A1 deletion leads to reduced incorporation of citrate into the TCA cycle. Moreover, our data support the hypothesis that mitochondria are a major source of citrate in the embryonic heart and that SLC25A1 deletion disrupts incorporation of citrate into the Krebs cycle thereby limiting oxidative metabolism.

To assess how SLC25A1 deletion impacts metabolic flux in the embryonic heart, we further integrated our experimental data with mathematical modeling using CardioNet ^40^, a genome-scale metabolic network of cardiac metabolism. To infer flux distributions in the embryonic heart, we used flux balance analysis and constrained internal metabolite fluxes based on our GC-MS-based metabolomics data (Fig. 5), as well as mitochondrial respiration rates (Fig. 4b). Linear programming was used to calculate solutions that satisfy the mass balance constraints and flux constraints. The objective function was defined as a maximization-minimization problem to reflect the unique requirements of the embryonic heart which maximizes ATP provision and biomass synthesis (e.g., protein and membrane lipid synthesis) while minimizing the utilization of extracellular substrates and oxygen from the plasma. Flux distributions were analyzed by PCA and among genotype-dependent separation, 80.6% of the total variance was captured in the first two dimensions (Fig. 6a). CardioNet simulations predict an increased utilization of exogenous glucose in response to SLC25A1 loss (Fig. 6b,c), which corresponds with increased glycolytic flux. However, glucose-derived pyruvate is entering the TCA cycle significantly less in response to loss of SLC25A1, which corresponds with a reduced TCA cycle flux and ATP provision (Supplemental Figure 6A, Fig. 6b), as well as reduced oxidative phosphorylation (Fig. 6d). Correspondingly, we identified a reduced glutamine-pyruvate transaminase flux compared to increased glutamic-oxaloacetic transaminase flux (Supplemental Figure 6B), suggesting that loss of SLC25A1 increases anaplerotic glutamine metabolism. Ultimately, as cardiac metabolism in the developing heart matures from dependence on glycolysis to incorporation of energy derived from mitochondrial oxidation, our simulations suggest that SLC25A1 loss impairs metabolic reprogramming and maturation in the developing heart.

**Figure 6.**
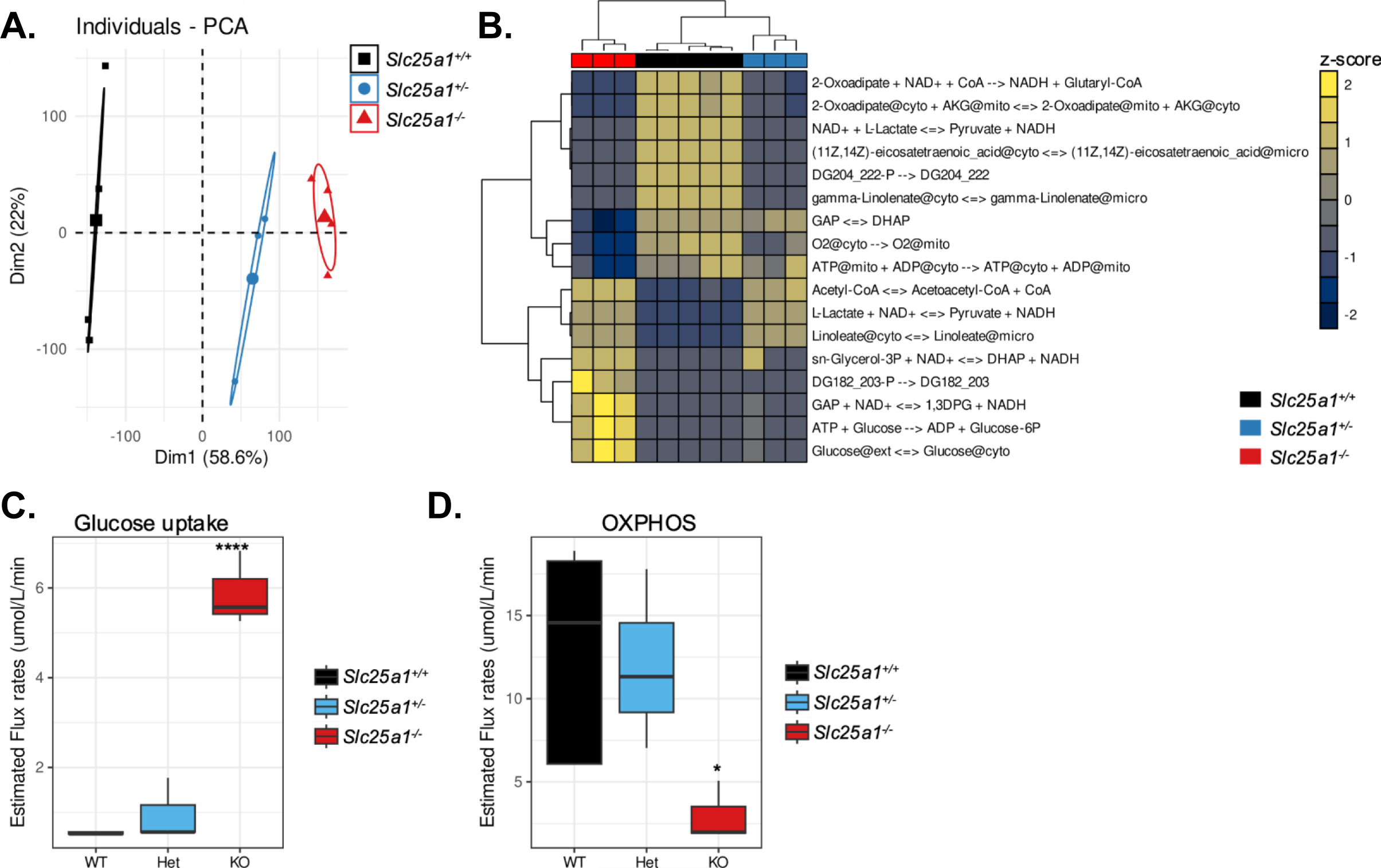
In silico modeling reveals glucose dependency and impaired oxidative phosphorylation with SLC25A1 loss. (A) PCA of the CardioNet simulation revealed a clear clustering of *Slc25a1^+/+^*, *Slc25a1^+/−^*, and *Slc25a1^−/−^* hearts by *Slc25a1* genotype. (B) Unsupervised hierarchical clustering of significantly altered flux distributions. Two-way ANOVA with post hoc Tukey’s test followed by multiple comparison analysis (FDR<1% after Benjamini, Krieger and Yekuteli). (C) Predicted flux rates for glucose uptake, and (D) oxidative phosphorylation in response to SLC25A1 deletion. *P < 0.05; ****P<0.00005

### *Slc25a1* deletion reduces histone acetylation in developing mouse hearts

Our combined transcriptomics, biochemical, and metabolomics analyses revealed defects in mitochondrial oxidative metabolism with *Slc25a1* deletion. We next asked what overarching regulatory mechanisms might control *Slc25a1* deletion-associated metabolic dysregulation. SLC25A1 is the main mitochondrial transporter responsible for citrate export ^10, 11^. In the cytosol, citrate is a precursor for synthesis of acetyl-CoA, a substrate for epigenetic modifications like histone acetylation ^17, 18^. As acetyl-CoA abundance modulates histone acetylation, SLC25A1 could connect mitochondrial metabolism to epigenetic regulation of gene expression via modulating citrate. To determine whether the observed decreased citrate levels in *Slc25a1^+/−^* and *Slc25a1^−/−^* hearts (Fig. 5d), alters histone acetylation, we assessed acetylation status of histone H3 at lysine 9 (H3K9), which is often enriched at promoters of actively transcribed genes ^41^, by immunoblots of total heart extracts from E17.5 *Slc25a1^+/+^, Slc25a1^+/−^*, and *Slc25a1^−/−^* embryos (Fig. 7a). We observed a reduced ratio of acetylated H3K9 (H3K9ac) to total histone H3 levels in *Slc25a1^+/−^* and *Slc25a1^−/−^* hearts compared to *Slc25a1^+/+^* controls (Fig. 7b), supporting a role for SLC25A1 in regulating histone acetylation in the developing heart.

**Figure 7.**
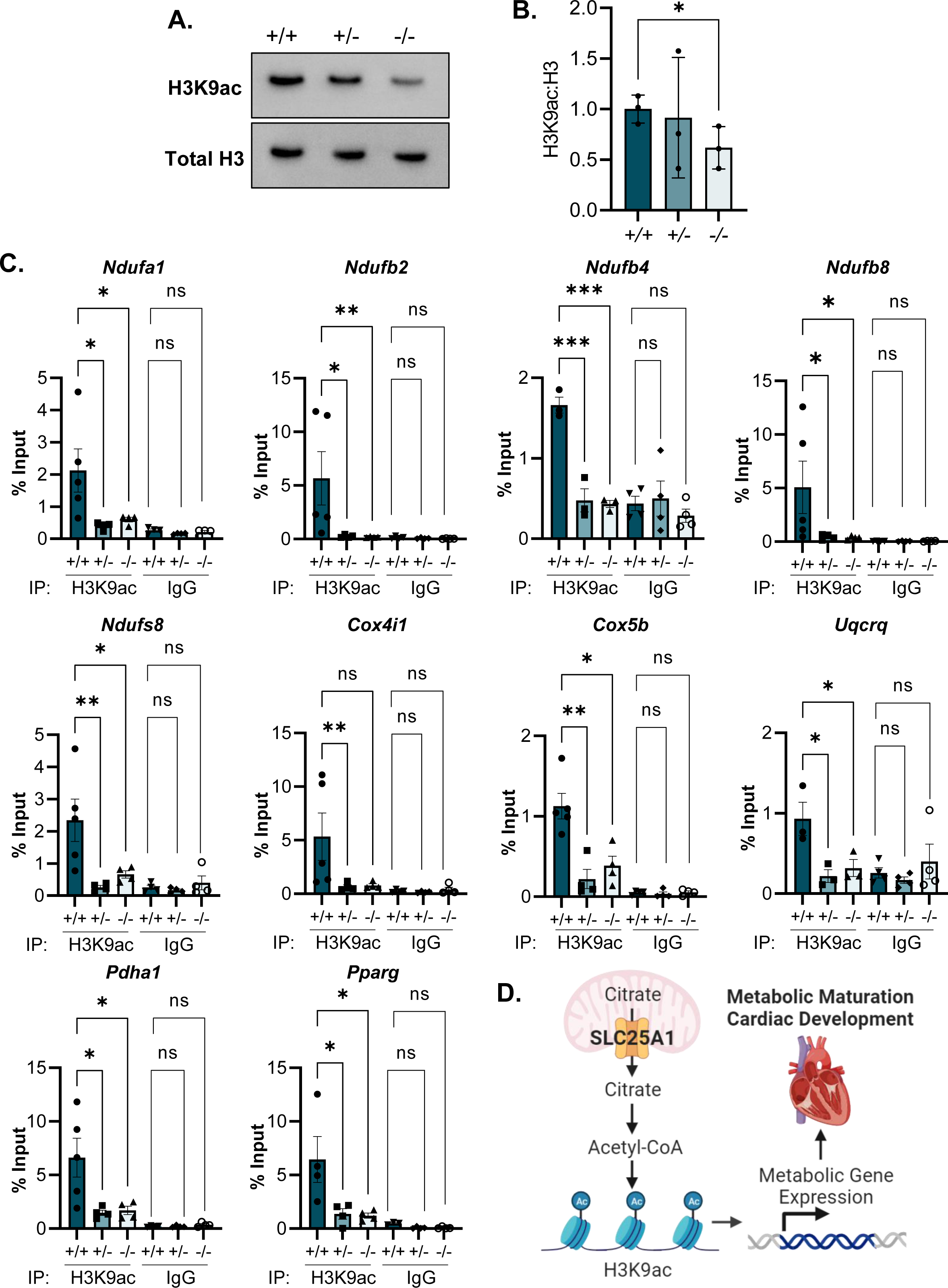
*Slc25a1* deletion reduces histone acetylation at promoter of metabolism-related genes in developing mouse hearts. (A) Representative western blot of histone H3 acetylated at K9 (H3K9ac) and total histone H3 in total heart protein from E17.5 embryos of the indicated genotypes. (B) Densitometry quantification of H3K9ac to H3 levels in E17.5 hearts (n = 3/group). One-way ANOVA followed by post hoc Dunnett’s test was used for statistical analysis. *P < 0.05. (C) H3K9ac ChIP qPCR of H3K9ac enrichment at the promoters of *Ndufa1*, *Ndufb2, Ndufb4*, *Ndufb8*, *Ndufs8*, *Cox4i1*, *Cox5b*, *Uqcrq*, *Pdha1* and *Pparg* in E17.5 hearts (n = 4-5/group). ChIP with anti-rabbit IgG was used as a negative control. Data were normalized using input DNA. Values presented as means ± SE. One-way ANOVA followed by post hoc Dunnett’s test was used for statistical analysis. ns denotes not significant; *P < 0.05, **P < 0.005, and ***P < 0.0005. (D) Proposed model of SLC25A1-mediated citrate export and its role in regulation of metabolic gene expression in the developing heart. Citrate in the mitochondria is generated from acetyl-CoA and oxaloacetate in the TCA cycle. SLC25A1 mediates mitochondrial citrate export. Citrate in the cytosol is converted to acetyl-CoA, which is used for histone acetylation (H3K9ac) and impacts gene expression necessary for proper heart development. Diagram created using BioRender.

We next postulated that SLC25A1-related histone acetylation changes could affect gene expression. Given technical challenges in profiling gene expression in large numbers of knockout embryonic hearts, we chose a candidate gene approach as a proof of concept. Because we observed a striking decrease in mitochondrial respiration in SLC25A1 deleted hearts, we chose to focus on genes associated with mitochondrial metabolism that were dysregulated in our Nanostring transcriptomics panel (*Ndufa1*, *Ndufb4*, *Ndufs8*, Ndufb2, *Ndufb8*, *Uqcrq*, *Cox4i1*, *Cox5b*, and *Pdha*) that are known to have H3K9ac enrichment at their promoters from analyses of publicly available H3K9ac chromatin immunoprecipitation-sequencing (ChIP-seq) datasets of the mouse heart ^41, 42^ (Supplemental Figure 7a). We additionally prioritized *Pparg* for analysis because it was also significantly downregulated in our transcriptomics analysis and has H3K9ac enrichment at its promoter (Supplemental Figure 7a). Importantly however, it is a well-characterized transcription factor that regulates expression of metabolism genes ^43^ and cardiomyocyte-specific deletion phenocopies the VSDs ^44^ we observe in SLC25A1 deleted embryos. Moreover, analysis of the SLC25A1-dependent metabolism gene set for potential shared transcription factor regulation by querying the TRRUST v2 database using Metascape identified this gene as a top potential transcription factor regulating the SLC25A1-dependent metabolism gene set (Supplemental Figure 7b). Thus, to test the possibility that SLC25A1 could impact the expression of these genes by regulating H3K9ac promoter enrichment, we performed ChIP for H3K9ac followed by quantitative PCR (qPCR) to assess H3K9ac abundance at the *Ndufa1*, *Ndufb4*, *Ndufs8*, *Ndufb2*, *Ndufb8*, *Uqcrq*, *Cox4i1*, *Cox5b*, *Pdha* and *Pparg* promoters in E17.5 *Slc25a1^+/+^, Slc25a1^+/−^*, and *Slc25a1^−/−^* embryonic hearts. Our results revealed significantly reduced H3K9ac at the promoters of *Ndufa1*, *Ndufb4*, *Ndufs8*, *Ndufb2*, *Ndufb8, Uqcrq*, *Cox5b*, *Pdha1* and *Pparg* in *Slc25a1^+/−^* and *Slc25a1^−/−^* hearts compared to *Slc25a1^+/+^* controls, as well as a trend for decreased H3K9ac deposition at the *Cox4i1* promoter (Fig. 7c). Collectively, these results support a role for SLC25A1 in gene expression regulation via histone acetylation in the developing heart.

## DISCUSSION

Though previous investigations have associated SLC25A1 with several diseases ^24, 26, 30–32, 35^, our results here provide comprehensive new evidence that SLC25A1 also contributes to metabolic reprogramming during cardiac development. Our mouse model results show that systemic loss of SLC25A1 led to stunted embryonic growth, perinatal lethality, and an array of developmental abnormalities including cardiac structural defects. Moreover, SLC25A1 loss resulted in mitochondrial dysfunction and profound perturbations in expression of genes associated with metabolism, as well as striking disruptions in cardiac mitochondrial ultrastructure and function that collectively point to a role for SLC25A1 in metabolic reprogramming during mouse heart development. Most importantly, *Slc25a1* haploinsufficiency was sufficient to produce cardiac developmental defects, which we believe underscores the clinical relevance of our finding of a near-significant association of ultrarare human *SLC25A1* variants associated with pediatric CHD.

Our initial efforts to identify the mechanisms by which *Slc25a1* deletion reduces respiratory chain function in the embryonic mouse heart led us to examine the relationship between SLC25A1 and the mitochondrial ribosome, as SLC25A1 expression regulates mitochondrial ribosome subunit expression in vitro ^21^. Our analyses of E17.5 hearts did not reveal any alterations in protein expression of key mitochondrial ribosome subunits with SLC25A1 loss, indicating that *Slc25a1* deletion-induced mitochondrial dysfunction is not due to impaired mitochondrial translation. As the mitochondrial proteome and function can differ between different tissues and cell types ^45, 46^, our data point to possible cell type- and tissue-specific effects of SLC25A1.

Integrating our mitochondrial functional studies, metabolomics and in silico studies to further understand the metabolic ramifications of SLC25A1 deletion confirmed dosage dependency of *Slc25a1* deletion-associated metabolic dysregulation and further highlighted a critical role of SLC25A1 in metabolic reprogramming in the developing mouse heart. Our analyses suggest that loss of SLC25A1 causes widespread dysregulation of cardiac metabolism, most significantly, by increasing glucose utilization and anaplerotic glutamine metabolism while impairing TCA cycle flux and oxidative metabolism. The immature developing heart is mainly fueled by glucose through glycolytic provision of ATP, while the maturing heart transitions to a metabolic omnivore which incorporates a wide-range of substrates including fatty acids, ketone bodies and amino acids. Our multi-omics analysis indicates that loss of SLC25A1 leads to a more ‘immature’ metabolic profile, whereby SLC25A1 deleted hearts remain reliant on glucose causing impaired oxidative metabolism.

How can the focal loss of a mitochondrial transporter protein such as SLC25A1 result in such widespread changes to the metabolic landscape of the developing heart? SLC25A1 is a mitochondrial inner membrane transporter that sits at the interface between the mitochondria and cytosol to export mitochondria-produced citrate to the cytosol ^14, 15^. In the mitochondria, as part of the TCA cycle, citrate is formed from condensation of acetyl-CoA and oxaloacetate and can be further oxidized to enable oxidative energy production ^47^. In the cytosol, citrate can be converted back into acetyl-CoA by ATP-citrate lyase ^48^. Acetyl-CoA is an essential substrate for many biological processes, including protein post-translational acetylation and epigenetic regulation of gene expression via histone acetylation and cellular acetyl-CoA levels has been shown to modulate histone acetylation status ^18, 48^. As acetyl-CoA is not freely permeable across the mitochondrial inner membrane, SLC25A1 may play a particularly important role in linking mitochondrial metabolism to epigenetic regulation of gene expression by modulating organelle compartmentalization of citrate and by extension, acetyl-CoA.

Our proof of concept ChIP studies of metabolism-related genes support a critical role for SLC25A1 in regulating gene expression in the developing heart through H3K9 acetylation (Fig. 7d). We found that SLC25A1 loss results in decreased total H3K9ac in the developing heart, leading to reduced H3K9ac abundance at the promoters of a number of mitochondrial metabolism-associated genes, which aligns with the reduced gene expression we observed in the transcriptomics panel and reduced mitochondrial respiration of SLC25A1 deleted hearts. Additionally, our finding of reduced H3K9ac at the PPARγ promoter was particularly notable, as embryos with cardiomyocyte-specific deletion of PPARγ phenocopy the VSDs we observed in mouse embryos with systemic *Slc25a1* deletion ^44^, further supporting a link between SLC25A1 and PPARγ in the developing heart. Collectively, our findings indicate that SLC25A1 is a mitochondrial regulator necessary for genetic reprogramming to establish oxidative metabolism in the maturing heart.

Our discovery that embryos with complete deletion of *Slc25a1* display cardiac malformations and mitochondrial dysfunction is notable but could be expected. However, our finding that *Slc25a1* haploinsufficiency in mice also produces cardiac malformations and mitochondrial dysfunction is striking and could help understand CHD. In humans, *SLC25A1* is located on chromosome 22q11.2 within the critical deletion region associated with 22q11.2DS^22^. Cardiac presentations of 22q11.DS commonly include conotruncal defects like VSDs ^23^, and additional congenital heart defects like left ventricular noncompaction can occur ^49, 50^. The cardiac defects associated with 22q11.2DS have largely been ascribed to TBX1, a transcription factor located in the critical deletion region that plays important roles in cardiovascular development ^51^. Yet, 22q11.2DS is a syndrome of haploinsufficiency and *Tbx1* haploinsufficiency does not reproduce the full spectrum of cardiac defects associated with 22q11.2DS; thus, additional factors must contribute to its cardiac phenotypes. Because partial loss of *Slc25a1* in mice is sufficient to produce CHD observed in 22q11.2DS, our data position *SLC25A1* as a novel 22q11.2 mitochondrial gene that contributes to the cardiac presentation of 22q11.2DS.

*SLC25A1* mutations underlie two rare autosomal recessive neurometabolic/neuromuscular disorders, DL-2HGA and CMS23 ^30–32^. While cardiac disease indices are not part of the OMIM characterization of these disorders, there have been hints that *SLC25A1* variants may also impact the heart. Indeed, VSDs, atrial septal defects, patent ductus arteriosus, bicommissural aortic valve, and patent formamen ovalae were reported in isolated cases of DL-2HGA due to *SLC25A1* mutations ^52–54^. Our analyses of the PCGC dataset revealed a near-significant (p = 0.058) association of ultrarare predicted damaging *SLC25A1* variants with pediatric CHD. While our *SLC25A1* variant burden testing did not reach statistical significance, it is important to note that our studies of the *Slc25a1* KO mouse model support neonatal lethality of *Slc25a1^+/−^* pups before P7. As such, predicted damaging *SLC25A1* variants may be underrepresented in the PCGC cohort due to lethality of undiagnosed neonates. Nevertheless, our analyses of the PCGC dataset strengthen the connection between *SLC25A1*, cardiac morphogenesis, and CHD, and support expansion of the clinical presentation of *SLC25A1* mutations to include cardiac structural defects. Moreover, we previously discovered that *SLC25A1* affects neurodevelopment in 22q11.2DS ^24^, while some DL-2HGA patients exhibit microcephaly ^52^, a feature we also observed in *Slc25a1^−/−^* mouse embryos. Thus, our work suggests that *SLC25A1* is a nuclear DNA-encoded mitochondrial gene that can connect the neurological and cardiac pathologies of 22q11.2DS and DL-2HGA.

In summary, we have uncovered a novel pathway of metabolic regulation in which SLC25A1 regulates gene expression in the developing heart. We demonstrated that *Slc25a1* is a dosage-dependent regulator of ventricular morphogenesis in the developing mouse embryo. Critically, our work in *Slc25a1* KO mice has clinical relevance, as we identified a strong trend for pathogenic variants of human *SLC25A1* to associate with CHD. Collectively, our work positions SLC25A1 as a mitochondrial transporter that mediates epigenetic regulation of gene expression in the developing heart and specifically highlights a role for SLC25A1 in metabolic maturation in heart development. As SLC25A1-dependent regulation of gene expression may extend beyond the confines of metabolism, our work also opens the door to a potential broader role for SLC25A1 in mitochondrial control of developmental programs. Finally, as *SLC25A1* is a 22q11.2 gene, and our combined mouse and human studies point to a role for *SLC25A1* as novel CHD risk factor, this study may be an important first step in defining SLC25A1 as a connection between cardiac and neurodevelopmental pathologies in 22q11.2DS, and the knockout mouse model offers an important tool for discovering the molecular basis of this condition.

## ACKNOWLEDGEMENTS

JQK was supported by the Additional Ventures Single Ventricle Research Fund, the Department of Defense (0000063651), and the National Institutes of Health (R01-GM-144729). VF was supported by the National Institutes of Health (1RF1AG060285). AK was supported by the National Institutes of Health (R00-HL-141702). GAP was supported by the National Institutes of Health (R01-HL144776). Transcriptomics for this study was supported in part by the Emory Integrated Genomics Core shared resource of Winship Cancer Institute of Emory University and the National Institutes of Health (NIH)/National Cancer Institute (P30CA138292). Electron microscopy for this study was supported by the Robert P. Apkarian Integrated Electron Microscopy Core, which is subsidized by the Emory University School of Medicine and the Emory College of Arts and Sciences. Additional support for electron microscopy was provided by the Georgia Clinical and Translational Science Alliance of NIH (UL1TR000454). Echocardiography for this study was supported by the Animal Physiology Core, which is subsidized by Emory University and Children’s Healthcare of Atlanta. Additional support was provided by the NIH Office of the Director (S10OD021748). Microscopy and imaging for this study was supported in part by the Emory University Integrated Cellular Imaging Core and Children’s Healthcare of Atlanta. The content is solely the responsibility of the authors and does not necessarily reflect the official views of funding sources.

## AUTHOR CONTRIBUTIONS

JQK and CO wrote the manuscript. JNP, CO, NG, AG, FS, JTG, MEB, and JQK performed experiments. AK conducted computational flux analysis and CardioNet modeling. JQK, VF, AK, MED, JNP, CO, JTG, and GAP analyzed the data. JQK, JNP, and CO designed the study. JQK conducted experimental oversight.

## SUPPLEMENTAL FIGURE LEGENDS

**Supplemental Table 1. Human *SLC25A1* variant KBAC burden test using pseudosibling controls.**

**Supplemental Figure 1. Expression SLC25A1 in the mouse embryonic myocardium and endocardium.** Representative immunofluorescence images of SLC25A1 expression in wild type embryo (E10.5, E12.5, and E14.5) and heart (E18.5) sections (6 μm). DAPI was used to counterstain nuclei, and MF20 was used to counterstain cardiomyocytes. Dashed yellow line outlines myocardium from the endocardium; asterisk indicates endocardium negative for SLC25A1.

**Supplemental Figure 2. S*l*c25a1 haploinsufficiency causes neonatal lethality in mice.** (A) Representative images of 6-μm longitudinal sections of *Slc25a1^+/+^ Slc25a1^+/−^* mouse hearts at 2 months of age stained with hematoxylin and eosin. (B) Heart weight to body weight ratios measured at 2 months of age (n = 13 *Slc25a1^+/+^*; n = 10 *Slc25a1^+/−^*). (C) Echocardiography measurements assessed in 2-month-old mice with the indicated genotypes (n = 6 *Slc25a1^+/+^*; n = 7 *Slc25a1^+/−^*). LVWAd denotes left ventricular anterior wall thickness at diastole, LVPWd denotes left ventricular posterior wall thickness at diastole, LVIDd denotes left ventricular interior dimension at diastole, EF denotes ejection fraction, FS denotes fractional shortening, SV denotes stroke volume, RVWT denotes right ventricular wall thickness, RVOT denotes right ventricular outflow tract, and TAPSE denotes tricuspid annular plane systolic excursion. Values reported as mean ± SEM. Student’s *t* test was used for statistical analysis. n.s. denotes not significant. (D) Frequency of genotypes of P7 pups obtained from crosses of *Slc25a1^+/+^* and *Slc25a1^+/−^* mice. Chi-squared test was used for statistical analysis. *P < 0.05.

**Supplemental Figure 3. Expression of key mitochondrial ribosome subunits are unchanged with SLC25A1 deletion.** Western blot of MRPS22 and MRPS18B from total heart protein from E17.5 embryos of the indicated genotypes. Coomassie staining of the gel was used as a protein loading control.

**Supplemental Figure 4. Quantification of SLC25A1 deletion associated mitochondrial ultrastructural defects in the developing heart.** Quantification of mitochondrial morphology to assess mitochondrial (A) area, (B) Feret’s diameter, (C) circularity, (D) aspect ratio, and (E) roundness (n = 3 *Slc25a1^+/+^*; n = 7 *Slc25a1^+/−^*; n = 3 *Slc25a1^−/−^* embryos analyzed; >200 mitochondria analyzed per genotype). Values reported as means ± SEM. One-way ANOVA followed by Dunnett’s post hoc test was used for statistical analysis. n.s. denotes not significant; ** P<0.005; ***P<0.0005; ****P<0.00005.

**Supplemental Figure 5. TOM20 levels are unchanged in SLC25A1-deleted hearts.**

(A) Western blot for TOM20 expression. Coomassie staining of the gel was used as a protein loading control. (B) Densitometry quantification of TOM20 expression (n = 3/group).

**Supplemental Figure 6. In silico modeling of TCA cycle, glutamine-pyruvate transaminase and glutamic-oxaloacetic transaminase flux with *Slc25a1* deletion.** Estimated flux rates of (A) TCA cycle and (B) glutamine-pyruvate transaminase and glutamic-oxaloacetic transaminase enzymes in *Slc25a1^+/+^*, *Slc25a1^+/−^*, *Slc25a1^−/−^* hearts as modeled using CardioNet. *P < 0.05; **P<0.005; ****P<0.00005

**Supplemental Figure 7.** H**3**K9ac **enrichment at promoter regions of dysregulated metabolic genes and transcription factors from Nanostring analysis.** H3K9ac ChIP-seq profiles for E12.5, E14.5, E16.5, and P0 mouse heart datasets (ENCSR968NPX, ENCSR345RKE, ENCSR373LNE, ENCSR519DNE) publicly available from the ENCODE portal ^41, 42^ were evaluated for enrichment at the target gene promoters.

## METHODS

### Animals

Slc25a1tm1a(EUCOMM)Wtsi KO-first mutant mice (*Slc25a1^+/−^*) were obtained from the Mutant Mouse Resource and Research Center at the University of California, Davis and maintained on a C57BL6/J background. Timed matings were established using 2–3-month-old *Slc25a1^+/−^* male and female mice, with the morning of an observed copulation plug established as E0.5. Experiments were conducted on embryos collected at E10.5, E12.5, E14.5, E17.5, and E18.5. Genotyping of *Slc25a1^+/−^* mice as well as embryos from *Slc25a1^+/−^* intercrosses was conducted on genomic DNA isolated from embryonic yolk sacs or tail biopsies. All animal experiments were carried out with experimental protocols reviewed and approved by Emory University’s Institutional Animal Care and Use Committee.

### Histology and immunostaining

Hearts were fixed in 10% formalin and embedded in paraffin. Paraffin-embedded tissues were sectioned (6 µm), deparaffinized, rehydrated, stained with hematoxylin and eosin (H&E), and imaged using a Nanozoomer 2.0-HT whole-slide scanner (Hamamatsu). Myocardial trabecular and compact layer thicknesses were measured on H&E-stained sections using QuPath image analysis software ^55^.

For immunofluorescence, paraffin-embedded sections were deparaffinized and rehydrated, and antigen retrieval was performed with citrate buffer (pH 6.0). The following primary antibodies were used: anti-myosin heavy chain (Developmental Studies Hybridoma Bank, MF20, 1:20), and anti-SLC25A1 (Proteintech, 15235-1-AP, 1:100). Secondary antibodies used were: donkey anti-mouse 488, and donkey anti-rabbit 594 (1:200, Cell Signaling Technologies). Slides were mounted with ProLong Gold Antifade Mountant with DAPI (Thermo Fisher Scientific) to counterstain nuclei. Fluorescence imaging was conducted using a Keyence BZ-X810 system.

### Whole exome sequencing and burden testing

Genomic analysis using the Pediatric Cardiac Genomic Consortium (PCGC) whole exome cohort followed published protocols ^56–58^. Briefly, whole-exome DNA from blood or salivary samples was captured using the Nimblegen v.2 exome capture reagent (Roche) or Nimblegen SeqxCap EZ MedExome Target Enrichment Kit (Roche), followed by Illumina DNA sequencing as previously described ^56–58^. The resulting sequencing data were processed at Yale University School of Medicine, and reads were mapped to the hg19 reference genome. Mapped reads were further processed using the GATK Best Practices workflows ^59^, as previously described ^56^. Single nucleotide variants and small indels were called with GATK HaplotypeCaller ^60^. Further data filtering was performed using plink ^61^, including removing individuals with low call rates, outlying heterozygosity rates, outlying relatedness rates, and sex discrepancy. A total of 3740 probands passed individual filtering, and there were a remaining 3401 complete trios. A case-control cohort was generated for all complete trios using pseudo-sibling controls. With this method, a proband’s genotypes are taken as the case, and a pseudo-control sample is created with all untransmitted alleles between the parents and the proband using plink ^62^. Variants were filtered on call rate, Hardy-Weinberg equilibrium, and a high number of Mendel errors (≥3). Variants within SLC25A1 were extracted using the longest known canonical hg19 SLC25A1 variant bed file downloaded from UCSC TableBrowser on 4/5/23 ^63^. The remaining variants were annotated using wANNOVAR ^64^. Loss of function (frameshift, splice site, start lost, stop gained, and stop loss) and missense variants with combined annotation-dependent depletion score >20 ^65^ and minor allele frequency <0.001 in all gnomAD ^66^ populations were kept for burden analysis. An additional variant set was created by keeping SLC25A1 variants that do not occur in the gnomAD exome database (v2.1.1, as of 4/12/23). Dichotomous traits were created by comparing probands with a specific phenotype to the rest of probands in the PCGC whole-exome cohort. Burden testing using these dichotomous traits was completed using the SKAT ^67^ package in R with the first three principal components as covariates. Burden testing using the pseudo-sibling-controlled cohort was performed using kernel based adaptive cluster methods using WISARD ^68^.

### Echocardiography

Mice were anesthetized with isoflurane (1.5%), and body temperature was maintained at 37°C. Echocardiography measurements were performed using a Vevo 2100 Imaging System (VisualSonics) equipped with a MS-400 transducer. M-mode and B-mode measurements were taken of the parasternal short- and long-axis views. Systolic and diastolic ventricular chamber dimensions as well as left and right ventricular wall thicknesses were assessed. Ejection fraction, fractional shortening, and stroke volume were calculated using VevoLab software.

### Metabolic gene expression profiling

Transcriptomic profiling of expression of 768 metabolism-related genes was conducted using the NanoString nCounter Mouse Metabolic Pathways Panel. Briefly, total RNA was isolated from E17.5 hearts (n = 4/genotype) using a RNeasy Micro Kit (Qiagen). NanoString processing was completed by the Emory Integrated Genomics Core. RNA quality was assessed with the NanoString protocol. mRNA counts were normalized by expression of the housekeeping gene *Tbp*. Data were analyzed using Qlucore Omics Explorer Version 3.7(24) or GSEA v4.3.2. Differentially expressed mRNAs from QLUCORE were analyzed with the Metascape engine to determine ontology enrichment as well as representation of transcription factors using TRRUST. GSEA analysis was used to query the REACTOME database ^69^.

### Transmission electron microscopy

Mouse heart samples were collected from E17.5 embryos and fixed in 2.5% glutaraldehyde in 0.1 M cacodylate buffer (pH 7.4). Hearts were subsequently postfixed in 1% osmium tetroxide and embedded in epoxy resin. Ultrathin sections (80–90 nm) were cut with a Leica EM CU6 microtome and counterstained with uranyl acetate and lead citrate. Sections were imaged on a JEOL JEM-1400 transmission electron microscope (Tokyo, Japan) equipped with a Gatan US1000 CCD camera (Pleasanton, CA). Mitochondrial morphology was quantified as previously described ^70^ using ImageJ software (NIH).

### Electrophoresis and immunoblotting

For western blots, total protein extracts were prepared from embryonic hearts solubilized in radioimmunoprecipitation assay buffer supplemented with combined protease/phosphatase inhibitor cocktail (Thermo Fisher Scientific). Proteins were resolved on 10% SDS-PAGE gels, transferred to PVDF membranes, immunodetected with antibodies, and imaged using a ChemiDoc system (BioRad). Primary antibodies used were: anti-SLC25A1 (Proteintech, 15235-1-AP, 1:1000), anti-TOM20 (Proteintech, 11802-1-AP, 1:1000), OXPHOS Blue Native WB Antibody Cocktail (contains antibodies against SDHA ATP5A, UQCRC2, NDUFA9, and COXIV; Abcam, ab110412, 1:500), anti-H3K9ac (Cell Signaling Technologies, 9649, 1:1000), anti-H3 (Cell Signaling Technologies, 4499, 1:1000), and anti-GAPDH (Fitzgerald, 10 R-G109A, 1:10,000). Secondary antibodies used were alkaline phosphatase-linked goat anti-rabbit IgG (Cell Signaling Technologies, 7054), alkaline phosphatase-linked goat anti-mouse IgG (Cell Signaling Technologies, 7056), horseradish peroxidase-linked goat anti-mouse IgG (Cell Signaling Technologies, 7076), and horseradish peroxidase-linked goat anti-rabbit IgG (Cell Signaling Technologies, 7074). Blots were imaged using a ChemiDoc XRS+ System (BioRad) and quantified using Image Lab software (BioRad).

### Mitochondrial oxygen consumption

Mitochondrial oxygen consumption was conducted on embryonic hearts as previously described^71^. Briefly, hearts were harvested from E17.5 embryos. Hearts were gently homogenized on ice in isolation buffer (225 mM mannitol, 10 mM sucrose, 0.5 mM EGTA, 10 mM HEPES, pH 7.4). The heart homogenate was diluted in respiration buffer (70 mM mannitol, 25 mM sucrose, 20 mM HEPES, 120 mM KCl, 5 mM KH_2_PO_4_, 3 mM MgCl_2_, pH 7.4) containing 5 mM glutamate and 3 mM malate. Oxygen consumption was measured using an Oxytherm System (Hansatech). Basal ADP-stimulated respiration was initiated with the addition of 1 mM ADP, uncoupled respiration was induced with the addition of 0.3 µM FCCP, and nonmitochondrial oxygen consumption was assessed with the addition of KCN. Basal and maximal uncoupled mitochondrial oxygen consumption rates were obtained by subtracting KCN-nonmitochondrial respiration, followed by normalization to protein concentration measured by Bradford assay (BioRad).

### mtDNA copy number analysis

Total DNA was prepared from snap-frozen embryonic heart tissue using a DNAeasy Kit (Qiagen). Four E17.5 hearts per genotype were pooled for each replicate (n = 4 replicates/genotype). Quantification of relative mtDNA copy number was conducted by measuring mtDNA/nDNA ratio using qPCR. Primers for mouse mtDNA (forward, CTAGAAACCCCGAAACCAAA; reverse, CCAGCTATCACCAAGCTCGT) and the β-2-microglobulin nuclear DNA (forward, 5’ ATGGGAAGCCGAACATACTG 3’; reverse, 5’ CAGTCTCAGTGGGGGTGAAT 3’) were used as previously described ^72^. qPCR was performed using LightCycler 480 SYBR Green I Master (Roche) on an Applied Biosystems QuantStudio 6 Flex Real-Time PCR system. Relative mtDNA copy number was calculated using the ΔΔC(t) method.

### Gas chromatography and mass spectrometry (GC/MS)-based targeted metabolomics

Metabolites were isolated as described previously ^73^. Briefly, frozen heart tissue sample were homogenized in 0.4 mL −80 °C cold MS-grade methanol, acetonitrile, water at a volume ratio of 40:40:20 (v/v/v/). Further disruption of tissue sample was achieved by freeze-thawing sample three-times. Extracts were centrifuged at 13,000 rpm for 5 min at 4 °C. The supernatant was transferred to clean glass tubes and D27-myristic acid (0.15 µg/µL, Cat No 366889, Sigma-Aldrich; St Louis, MO) was added as internal run standard. Vials were covered with a breathable membrane and metabolites were evaporated to dryness under vacuum. Dried samples were stored at −80°C for further GC-MS analysis. For derivatization of polar metabolites frozen and dried metabolites from cell extractions were dissolved in 20 µL of Methoxamine (MOX) Reagent (2% solution of methoxyamine-hydrogen chloride in pyridine; Cat. No. 89803 and 270970; Sigma-Aldrich; St Louis, MO) and incubated at 30°C for 90 minutes. Subsequently, 90 µL of N-tert-Butyldimethylsilyl-N-methyltrifluoroacetamide with 1% tert-Butyldimethyl-chlorosilane (TBDMS; Cat. No. 375934, Sigma-Aldrich; St Louis, MO.) was added and samples were incubated under shaking at 37°C for 30 minutes. GC-MS analysis was conducted using an Agilent 8890 GC coupled with a 5977-mass selective detector. Metabolites were separated on an Agilent HP-5MS Ultra Inert capillary column (Cat.No. 19091S-433UI). For each sample, 1 µL was injected at 250°C using helium gas as a carrier with a flow rate of 1.1064 mL/min. For the measurement of polar metabolites, the GC oven temperature was kept at 60°C and increased to 325°C at a rate of 10°C/min (10 min hold) followed by a postrun temperature at 325°C for 1 min. The total run time was 37 min. The MS source and quadrupole were kept at 230°C and 150°C, respectively, and the detector was run in scanning mode, recording ion abundance within 50 to 650 m/z.

### Mathematical modeling of the myocardial metabolic adaptations in the embryonal heart

*In silico* simulations were performed using the metabolic network of the cardiomyocyte CardioNet ^40, 73, 74^. Mathematical modeling can accurately predict the dynamics of cardiac metabolism in response to stress, and CardioNet can identify metabolic profiles and flux distributions. Flux balance analysis (FBA) allows to estimate flux rates by integrating experimental constraints on metabolites or proteins within a defined biochemical network. We specified flux constraints for extracellular metabolites reflecting the circulating metabolite composition in fetal plasma. Intracellular metabolites were constrained using experimentally determined metabolite abundance (GC-MS). Based on these constraints we first determined flux distributions (*v*_*m*_) under *Slc25a1^+/+^* (control) conditions. We then calculated fold-changes (FC) for experimentally measured metabolite concentrations between control and experimental groups (*Slc25a1^+-+^ and Slc25a1^−/−^*) and used these fold-changes to further constrain fluxes (*v*_*m*_) for the synthesis and/or degradation of intracellular metabolites. We included fold-changes (FC) based on the assumption that changes in metabolite concentrations under experimental conditions are accompanied by a proportional increase or decrease in the respective flux for the metabolite pool. By using metabolite level changes (fold changes) to estimate flux rate changes (*v_FC_*), we imply that the altered steady-state concentrations of metabolites are reflected in the newly evolved flux state and potentially limit metabolic functions.

To reflect the unique growth requirements in the fetal heart, we defined a biomass function including the synthesis of structural proteins, membrane lipids and ATPase activity ^40^. Linear problems were defined for experimental group to identify steady-state flux distributions that fulfill the applied constraints (e.g., substrate uptake and release rates), as well as changes in the metabolite pools. Regular FBA seeks to minimize the sum of all internal fluxes to fulfill applied constraints and the steady conditions ^40^. However, in the context of a growing heart this condition may not yield correct flux estimations, because the maximization of biomass is not reflected in a regular minimization optimization problem. Instead, we defined the objective as a maximin problem, which maximizes the minimum objective (e.g., flux) for all a given set of internal constraints ^40^

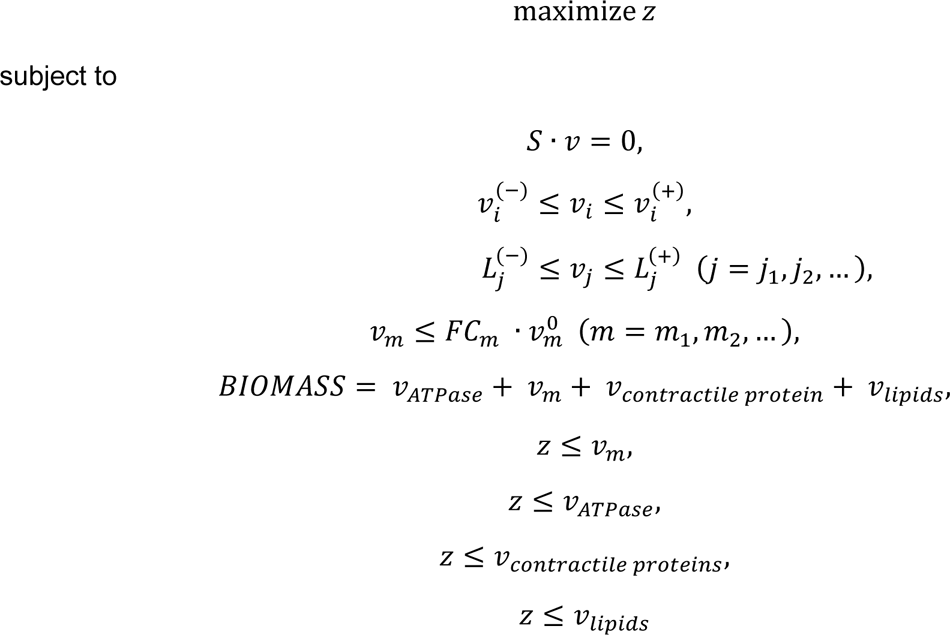

where *v*_*i*_ denotes the flux rate change through reaction *i*, *v*_*j*_ denotes the measured uptake or secretion rate through reaction *j*, *S* is the stoichiometric matrix, *v*_*i*_^(−)^ and 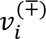 are flux constraints, and *z* that is an upper bound for each of the individual variables within the biomass function. Linear problems were implemented in perl and solved using the GUROBI LP solver ^75^ The resulting fluxes were compared between groups using a two-way ANOVA followed by multiple comparisons analysis. Adjusted q-values were computed using the Bonferroni, Krieger and Yekuteli two-step method.

### ChIP-qPCR and analyses of ChIP-seq data

ChIP experiments were conducted using the Magna ChIP HiSens Chromatin Immunoprecipitation Kit (EMD Millipore) with modifications. Briefly, E17.5 hearts were collected and pulverized with a mortar and pestle in liquid nitrogen. DNA–protein interactions were preserved with crosslinking buffer at continuous rotation for 20 minutes. The crosslinking reaction was stopped with addition of 2.5 M glycine. Following centrifugation (2000 *g* for 15 minutes) and washing with cold 1X PBS, samples were centrifuged again (2500 *g* for 10 minutes). Nuclei isolation buffer and sonication/ChIP/wash buffer were added to each sample for cell lysis and release of crosslinked protein–DNA. Samples were sonicated on ice with Misonix XL-2000 output 3 for 50 cycles of 15 seconds with a 45-second refractory period. ChIP was performed according to the manufacturer’s instructions. Antibodies used were: anti-H3K9ac (Sigma Aldrich, 17-658), and normal rabbit IgG (Sigma Aldrich, 12-370). Immunoprecipitated and input DNA was subjected to qPCR on a CFX96 Touch Real-Time PCR Detection System (BioRad) using SsoAdvanced Universal SYBR Green Supermix (Bio Rad 172-5270). The following primer sets were used to amplify targeted promoter regions: *Pparg* forward, 5’ CTGCGTAACTGACAGCCTAACC 3’ reverse, 5’ CTTGTGTGACTTCTCCTCAGCC 3’; *Ndufa1* forward, 5’ CCCCGTTGGTGAATTTGTGG 3’ reverse, 5’ GCGGAGATGTGGTTCGAGAT 3’; *Pdha1* forward, 5’ AGAACTAGGCCCTCAGACGA 3’, reverse, 5’CATGAGGAAGATGCTTGC CG 3’; *Ndufs8* forward, 5’ AACCTGCAGAGTGACCTTGG 3’, reverse,5’ TTGAACCGCATC TGGGCC 3’; *Cox4i1* forward, 5’ TCTGAAACGGACGGGCTTG 3’, reverse,5’ GACCGCCCTCTA CACCCT 3’; *Ndufb2* forward, 5’ CGGGACTCGGGGAAGTGA 3’, reverse,5’ CTGGACCCGTCT TGTCTCTG 3’; *Ndufb8* forward, 5’ CACTCACCCCATGCCATCAT 3’, reverse,5’ CAGAGTGA ACCGCGGAGAAG 3’; *Cox5,* forward, 5’ GCGTTGTTAGACTCCCACCA 3’, reverse, 5’CCACT CCGCGAAGTAACCTT 3’; *Ndufb4* forward, 5’ GGGGTCGTTGTACTGAAGCA3’, reverse,5’ GC ACTTCCTGAGCCTGAAGG 3’; *Uqcrq* forward, 5’ GTCTTTCAGAGGCAGGGGAC 3’, reverse,5’ GCAACCAGGAGTTTGAGCAG 3’. Percent input was calculated using the ΔΔC(t) method ^76^.

