## Supplementary figures and images for "Mitochondrial citrate carrier SLC25A1 is a dosage-dependent regulator of metabolic reprogramming and morphogenesis in the developing heart"

### Supplemental Figure 1

**Supplemental Figure 1.**

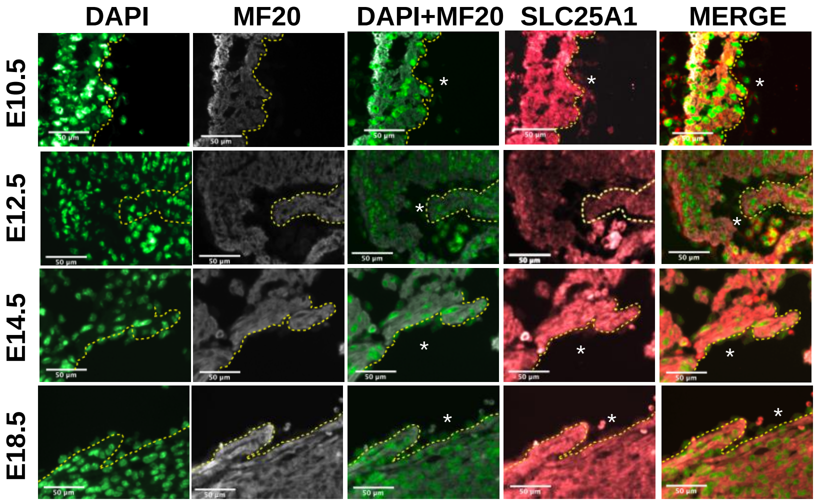

### Supplemental Figure 2

**Supplemental Figure 2.**

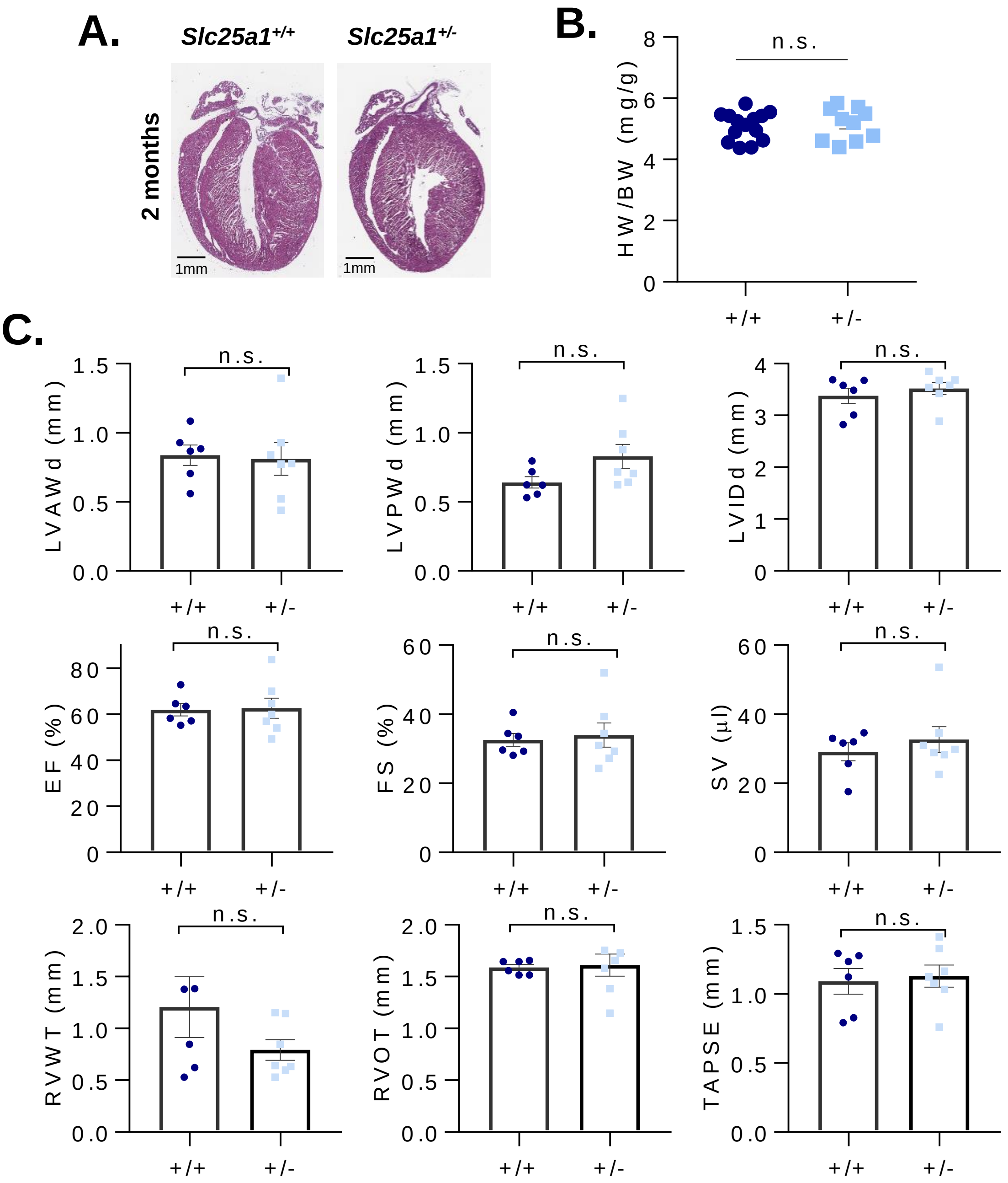


D.

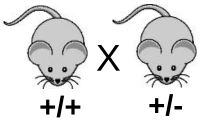


| | Genotype: | |
|---|---|---|
| Age: | +/+ | +/- |
| P7 | 56% (50%) | 44%* (50%) |

*Observed* / *Expected*

### Supplemental Figure 3

**Supplemental Figure 3.**

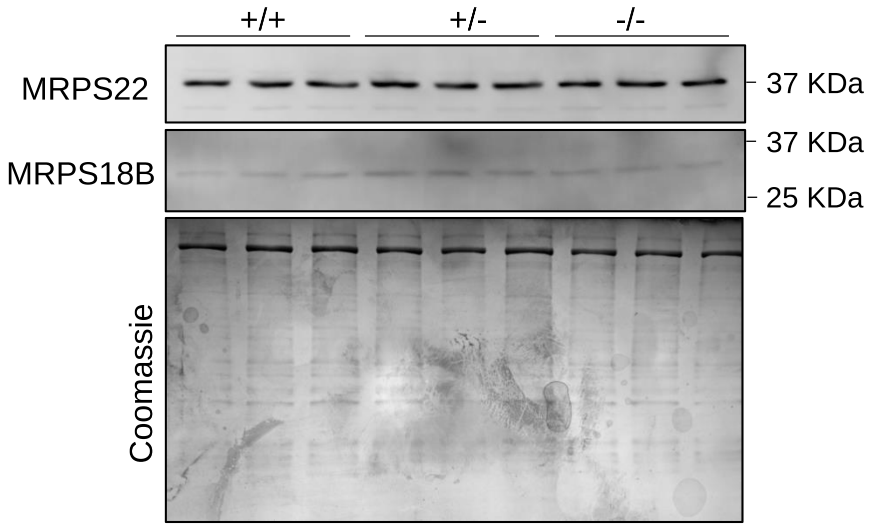

### Supplemental Figure 4

**Supplemental Figure 4.**

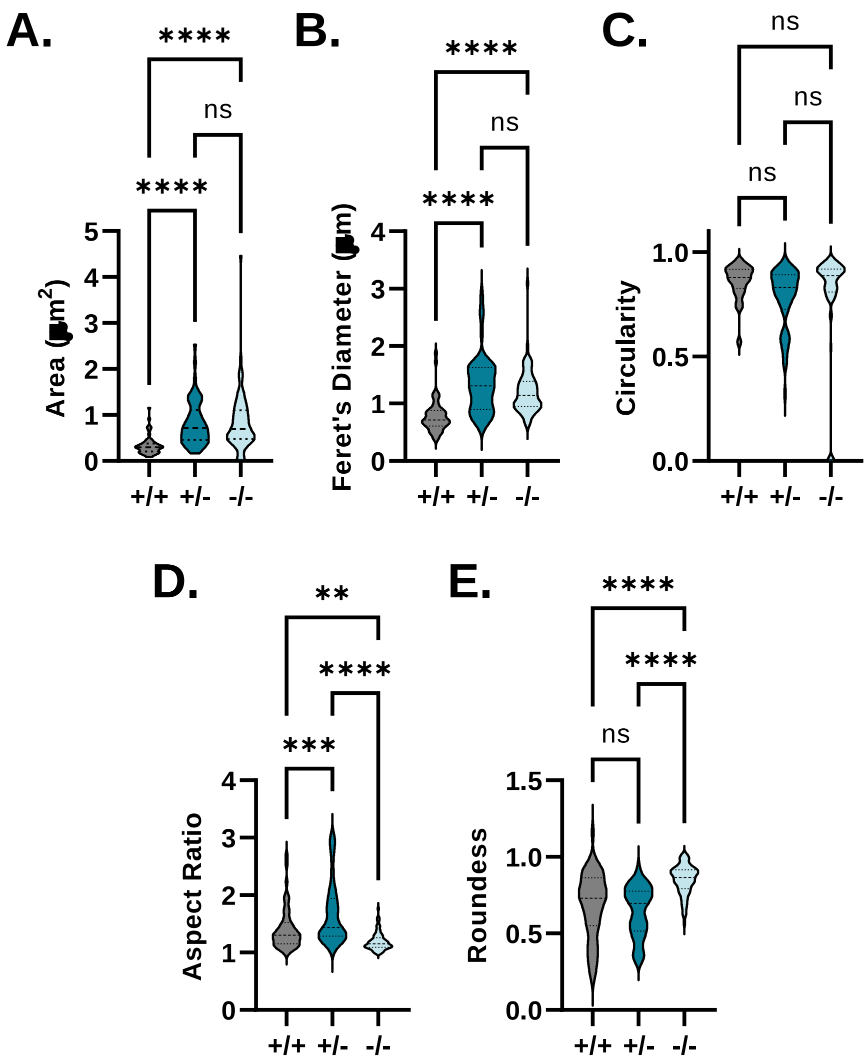

### Supplemental Figure 5

**Supplemental Figure 5.**

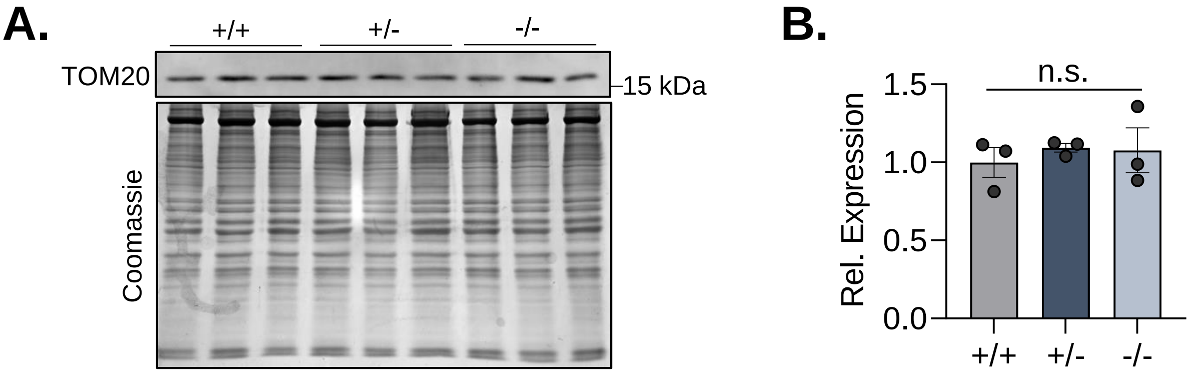

### Supplemental Figure 6

**Supplemental Figure 6.**

# A.

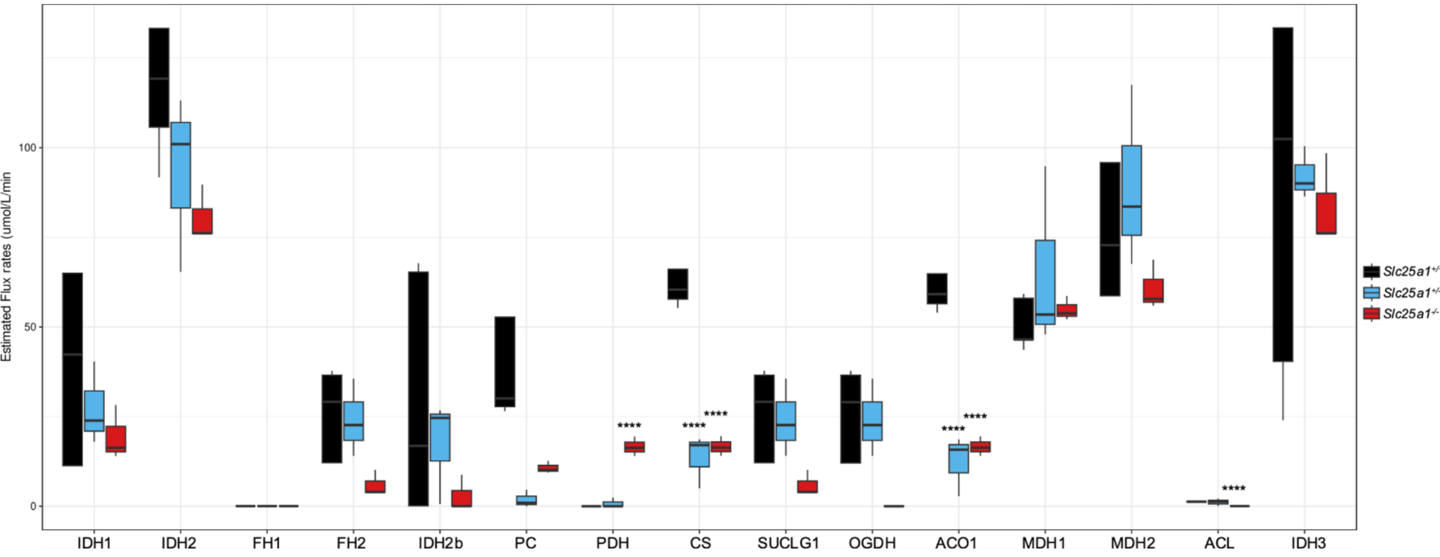


# B.

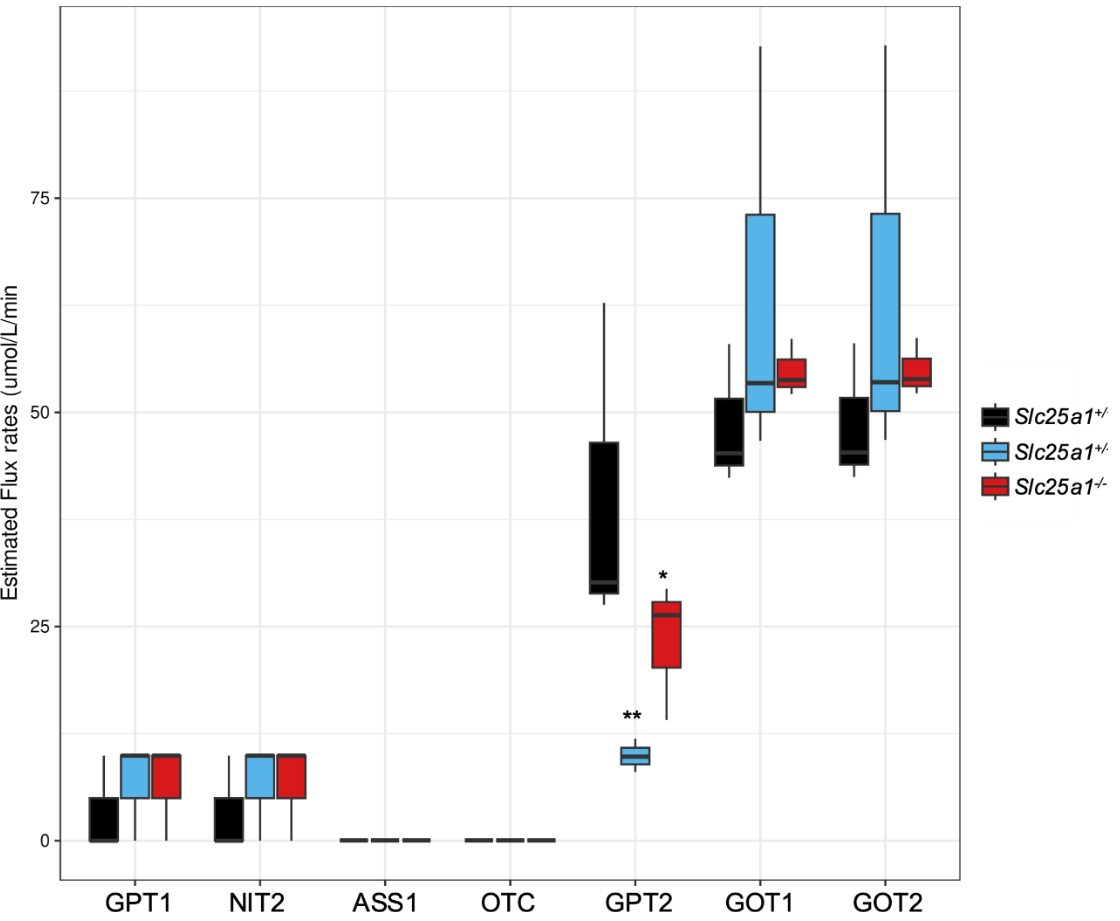

### Supplemental Figure 7

#### Supplemental Figure 7.

**A.**

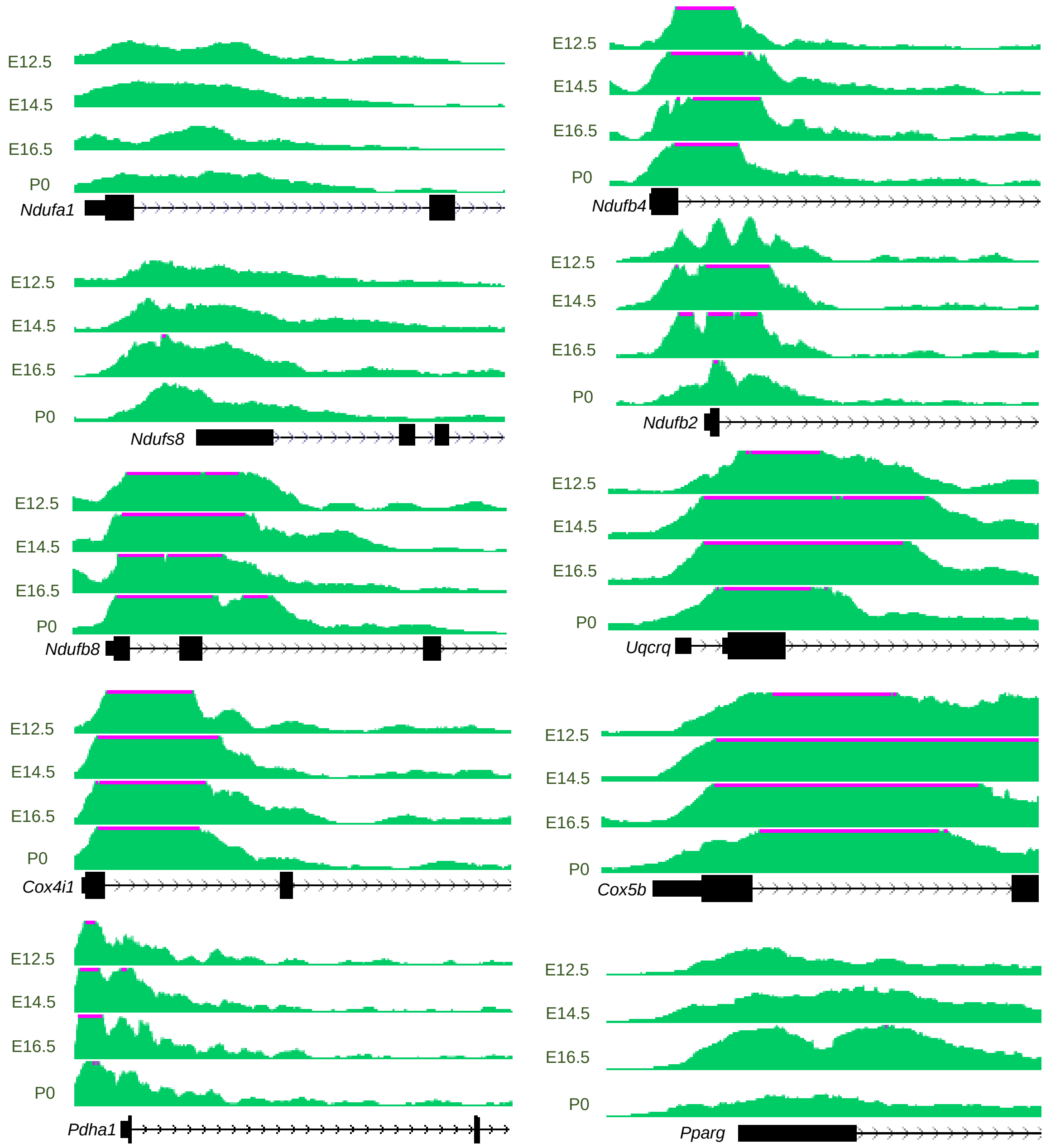


**B.**

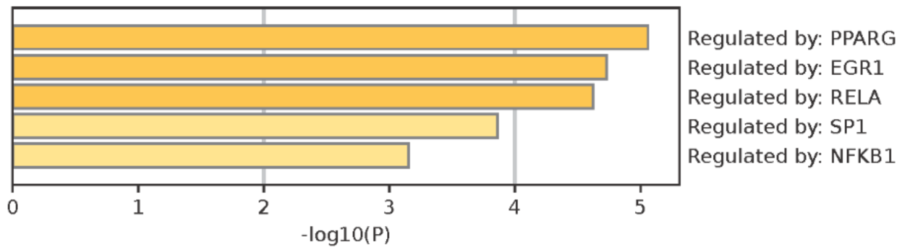
