## Supplemental Table 1 for "Mitochondrial citrate carrier SLC25A1 is a dosage-dependent regulator of metabolic reprogramming and morphogenesis in the developing heart"

**Table 1. Human *SLC25A1* Variant KBAC Burden Test using Pseudosibling Controls.**

| Variant Types | Variant N | KBAC p-value |
|---|---|---|
| **Gene Wide Nonsynonymous** | 25 | 0.61039 |
| **Gene Wide Nonsynonymous, CADD >20** | 17 | 0.48951 |
| **All ultrarare** | 34 | 0.91109 |
| **Ultrarare intronic** | 19 | 1.00000 |
| **Ultrarare nonsynonymous/ stop-gain** | 12 | 0.05794 |
